## Supplementary Figures for "Proteogenomic Discovery of Neoantigens Facilitates Personalized Multi-antigen Targeted T cell Immunotherapy for Brain Tumors"

**Supplementary Figures 1-19**

**a**

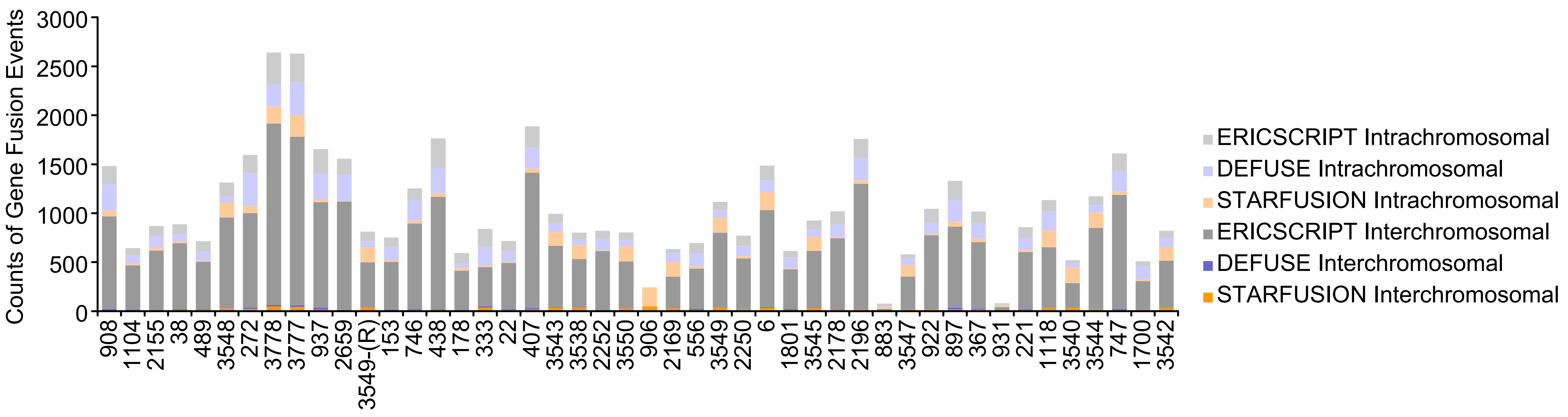


**b**

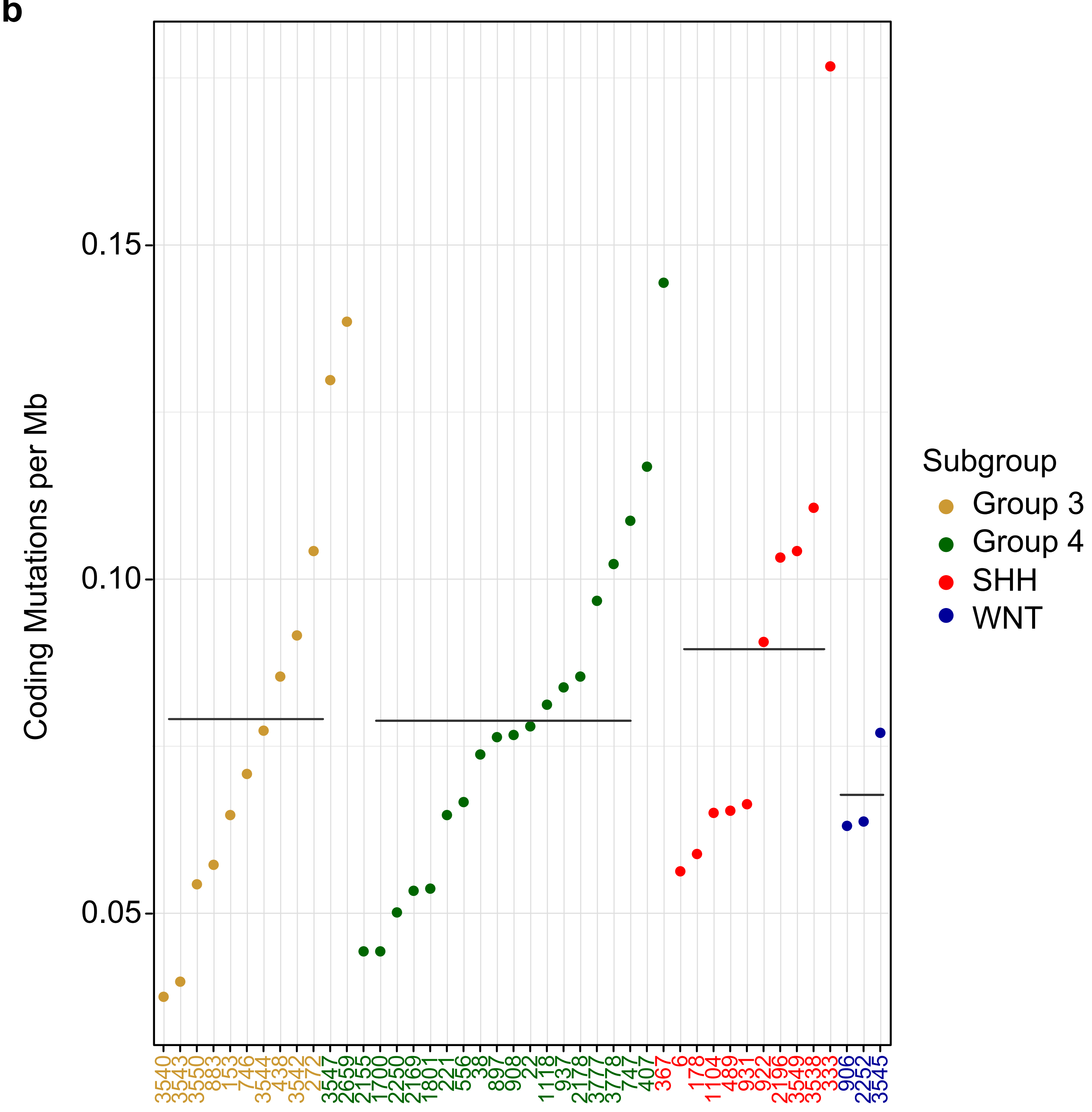


**Supplementary Fig. 1**

**Supplementary Fig. 1. Characterization of the gene fusion and somatic coding mutation events identified in medulloblastoma tumors. a** Aggregate barplot indicating the number of fusions detected in medulloblastoma tumors. Inter and intra chromosome fusions for each fusion caller are represented. **b** Plot indicating the number somatic coding mutations per megabase identified in the medulloblastoma tumors. The mean is indicated as a horizontal line.

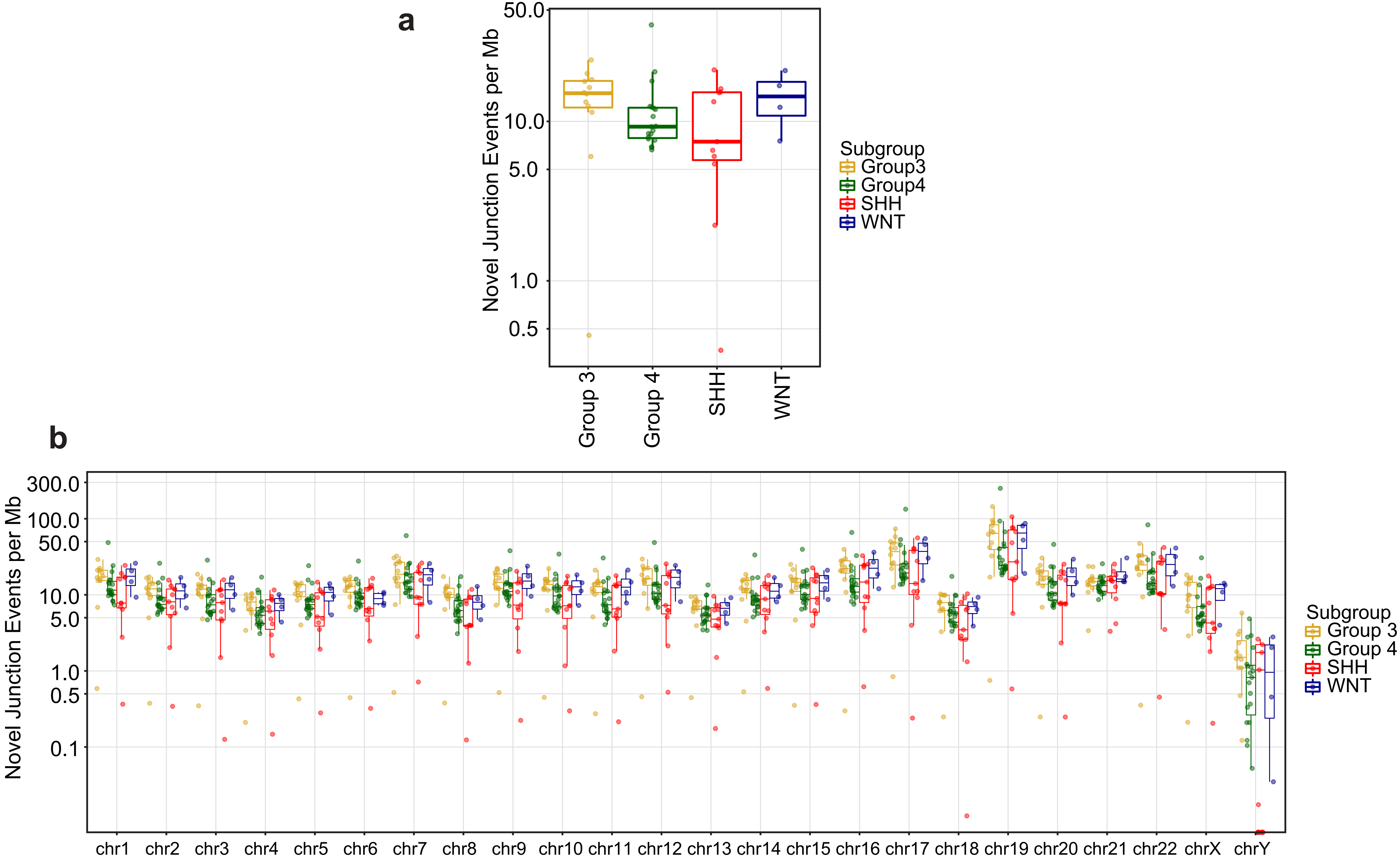


**Supplementary Fig. 2**

**Supplementary Fig. 2. Novel junction events identified in medulloblastoma tumors.** Boxplot indicating the number of novel junctions per medulloblastoma subgroup (**a**) or per chromosome identified in the medulloblastoma tumors (**b**). In the box plots, the lower and upper hinges correspond to the first and third quartiles, the middle line indicates the median. The upper whisker extends from the hinge to the largest value no further than 1.5 times of the interquartile range (IQR). The lower whisker extends from the hinge to the smallest value at most 1.5 times of the IQR.

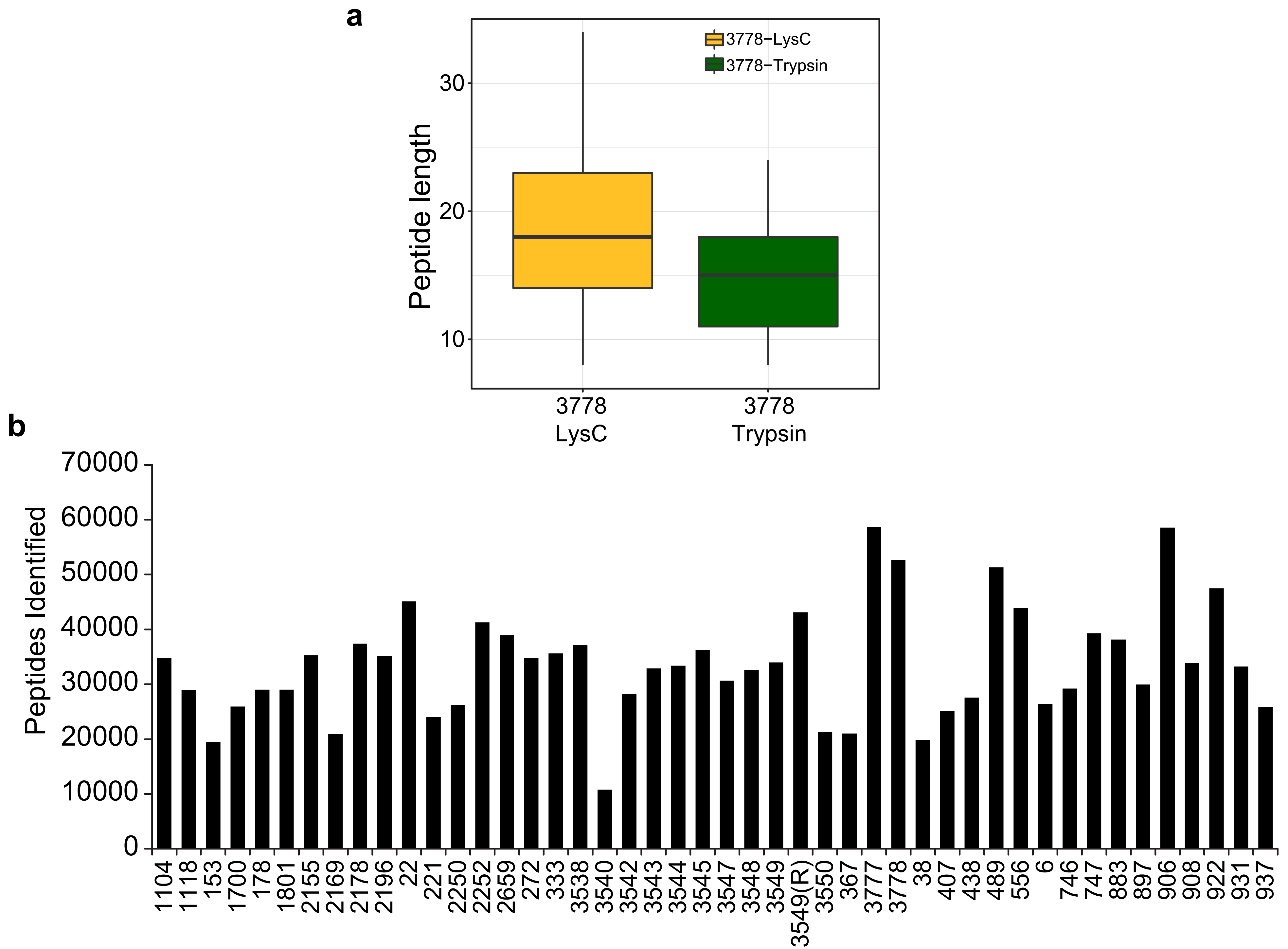


**Supplementary Fig. 3**

**Supplementary Fig. 3. Summary of proteomic findings in 46 medulloblastoma tumors. a** Boxplot representing the length of the peptides identified in the 7316-3778 patient by LC-MS/MS using trypsin or LysC enzymes. **b** Total number of unique peptides identified by LC-MS/MS in 46 medulloblastoma tumor tissues. In the box plot, the lower and upper hinges correspond to the first and third quartiles, the middle line indicates the median. The upper whisker extends from the hinge to the largest value no further than 1.5 times of the interquartile range (IQR). The lower whisker extends from the hinge to the smallest value at most 1.5 times of the IQR.

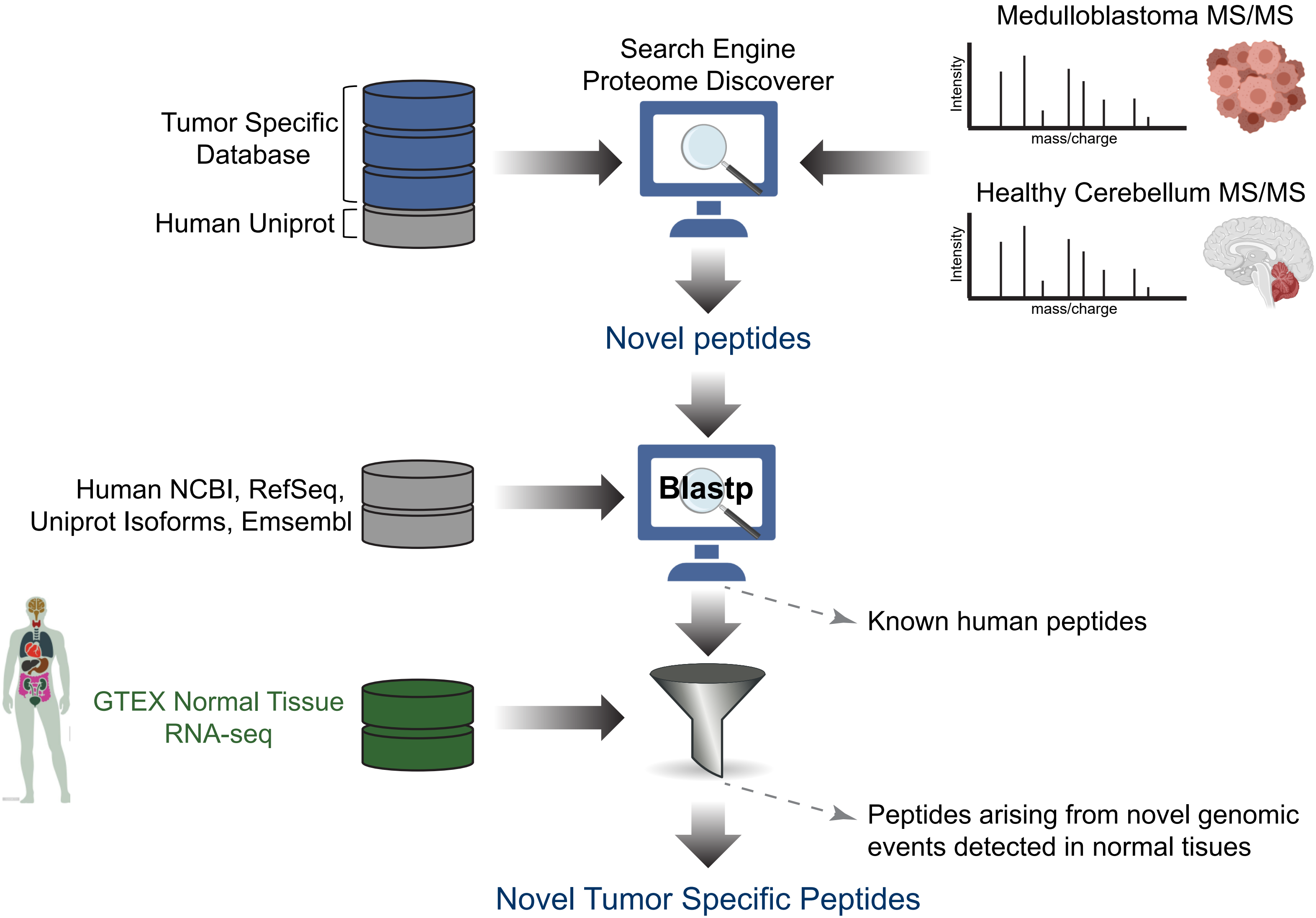


**Supplementary Fig. 4**

**Supplementary Fig. 4. Tumor specific peptide filtering strategy.** Cell lines and medulloblastoma tissues were subjected to LC-MS/MS shotgun proteomics and spectra were searched against tumor-specific databases generated from tumor WGS and RNA-seq. LC-MS/MS data from age matched healthy cerebella were included to identify unannotated peptides in healthy cerebellar tissues. The identified tumor specific peptides were BLASTed to human proteins from different databases (NCBI, RefSeq, Uniprot, neXProt, and Ensembl) to remove annotated peptides. Finally, to remove unannotated peptides in normal tissues other than cerebellum, databases were constructed using healthy tissue from different organs using RNA-seq from the GTEX project. Peptides arising from novel genomic events identified in normal tissues were removed.

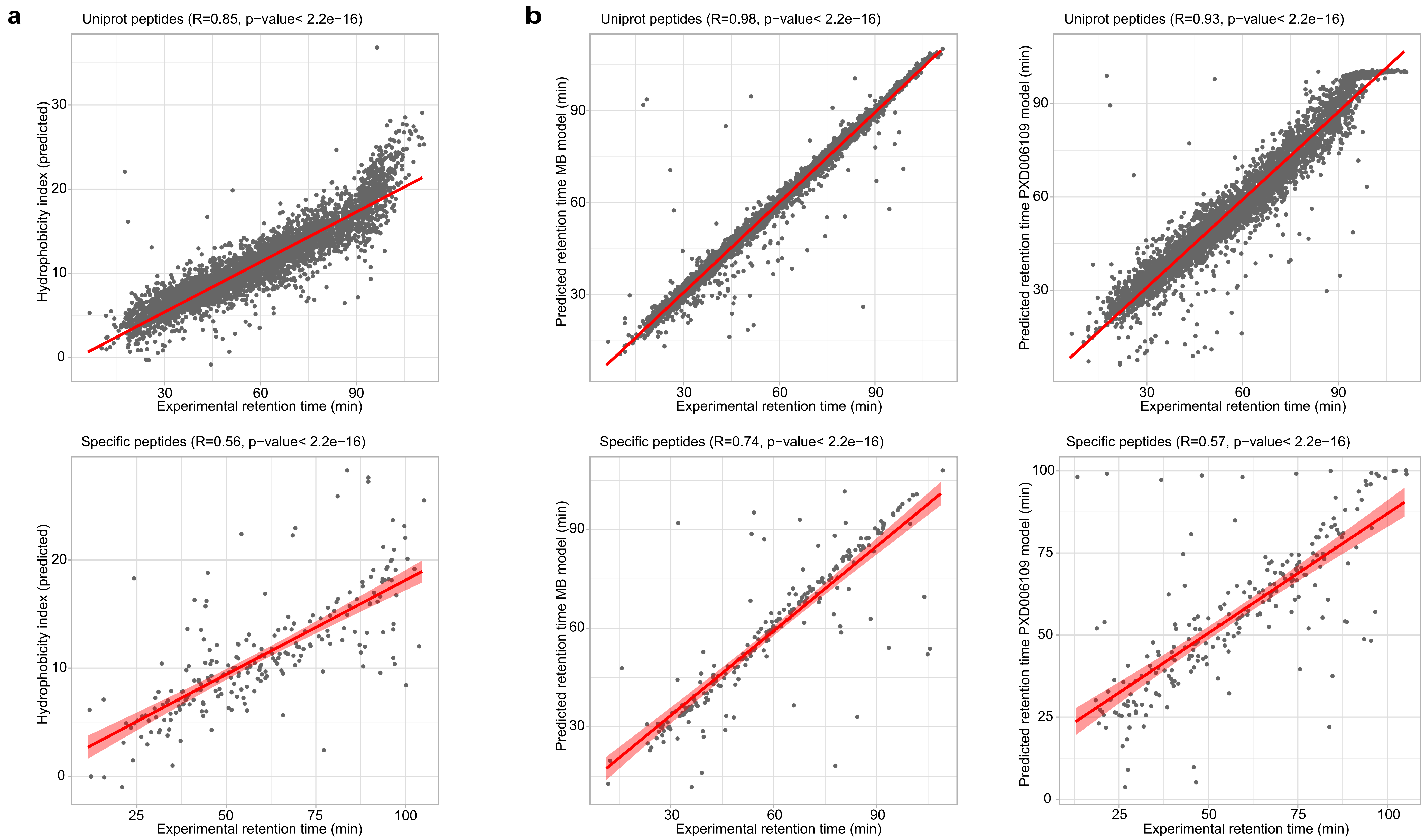


**Supplementary Fig. 5**

**Supplementary Fig. 5. Retention time validation of the identified peptides in medulloblastoma tumors**. **a** The hydrophobicity index for peptides identified in the medulloblastoma tumors were plotted against the observed retention time. Uniprot peptides are displayed in the upper row, medulloblastoma specific peptides are represented in the lower row. **b** Peptide retention times were predicted with the AutoRT algorithm using 2 different models: MB models using all peptides identified in this study (left) and the PXD006109 model (right). Predicted retention times were potted against experimental retention times. Uniprot peptides are displayed in the upper row, medulloblastoma specific peptides are represented in the lower row. R squared and p-values were calculated fitting the data to a linear regression model using the R function lm; p-values are indicated in the figure.

Peptide Validation 7316-3778

AGGAADMTDNIPLQPVR

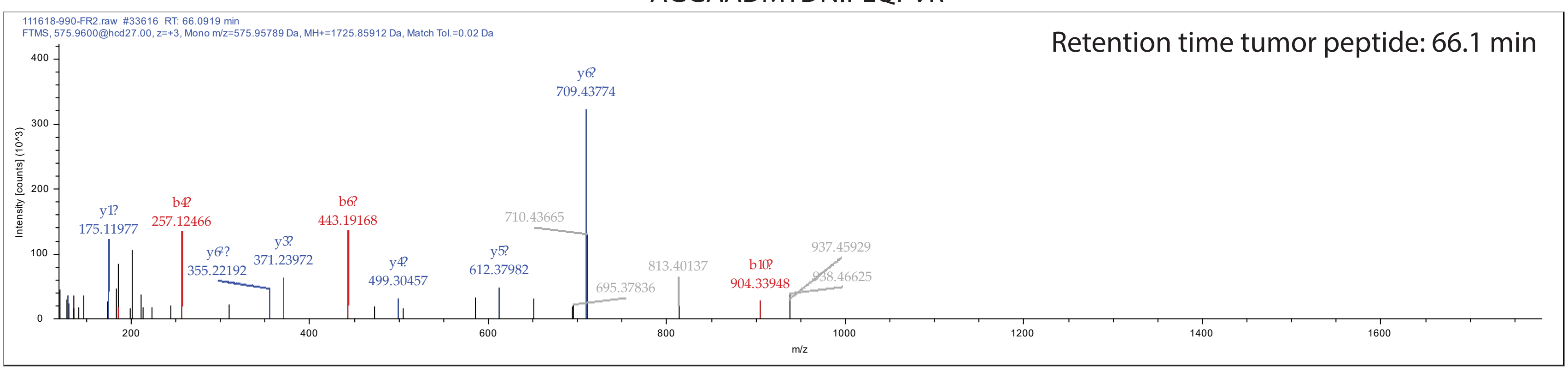


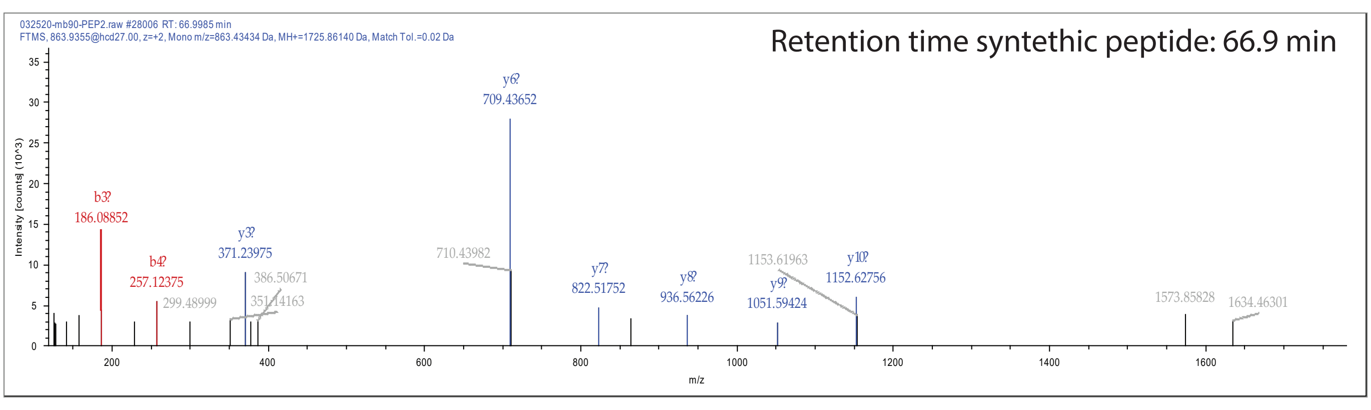


AGGAADMTDNIPLQPVRQK

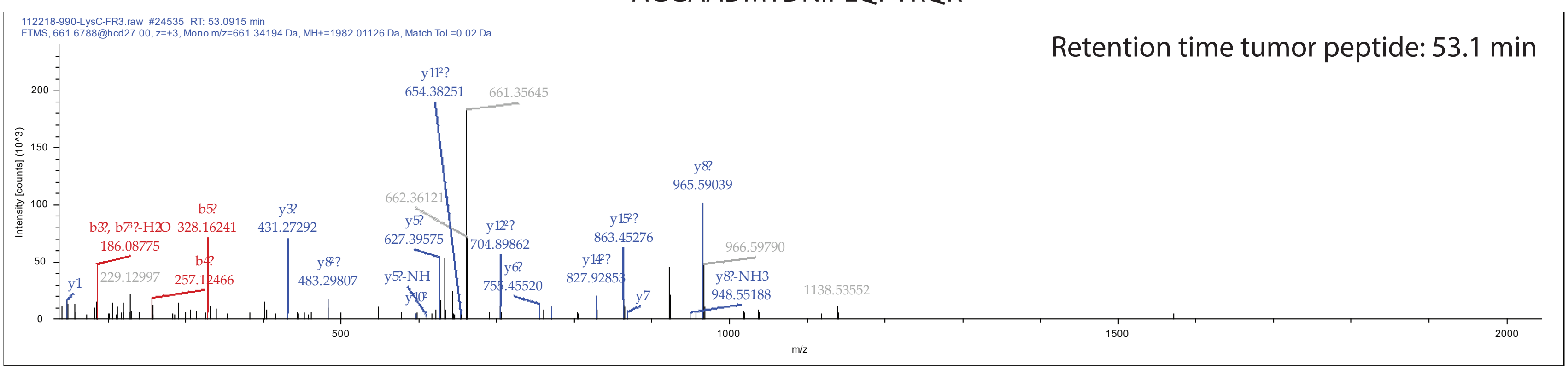


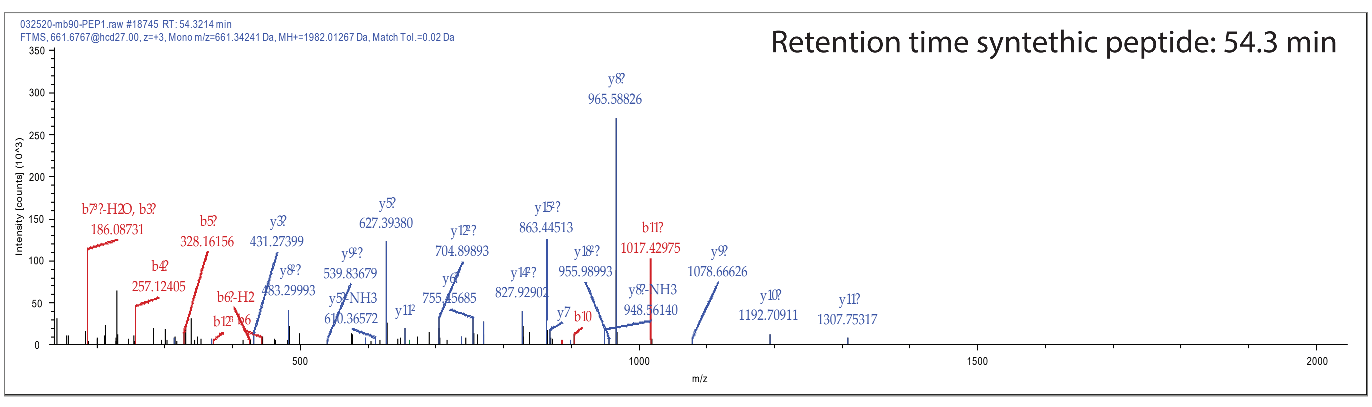


INEVLTSSPSPK

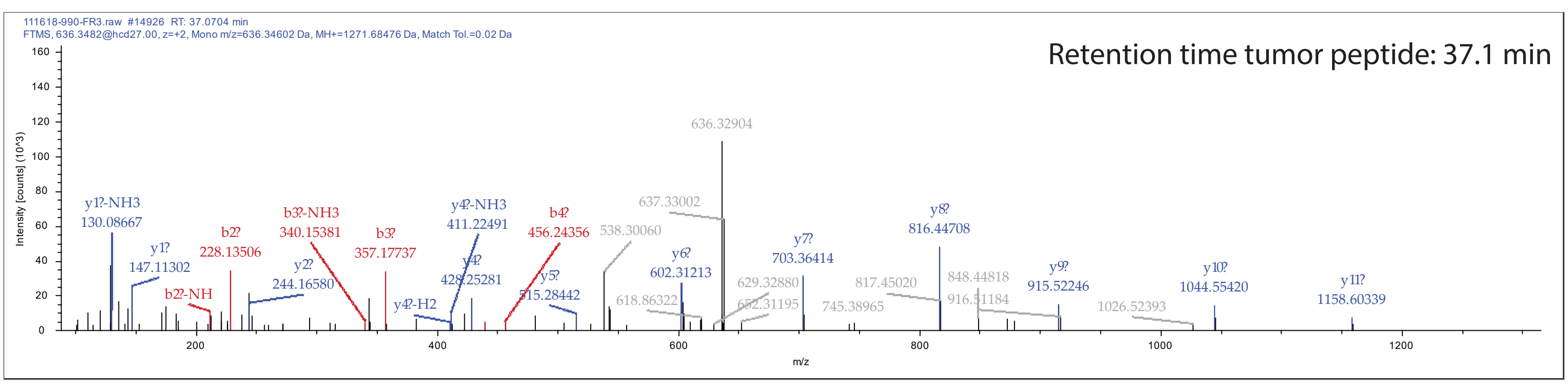


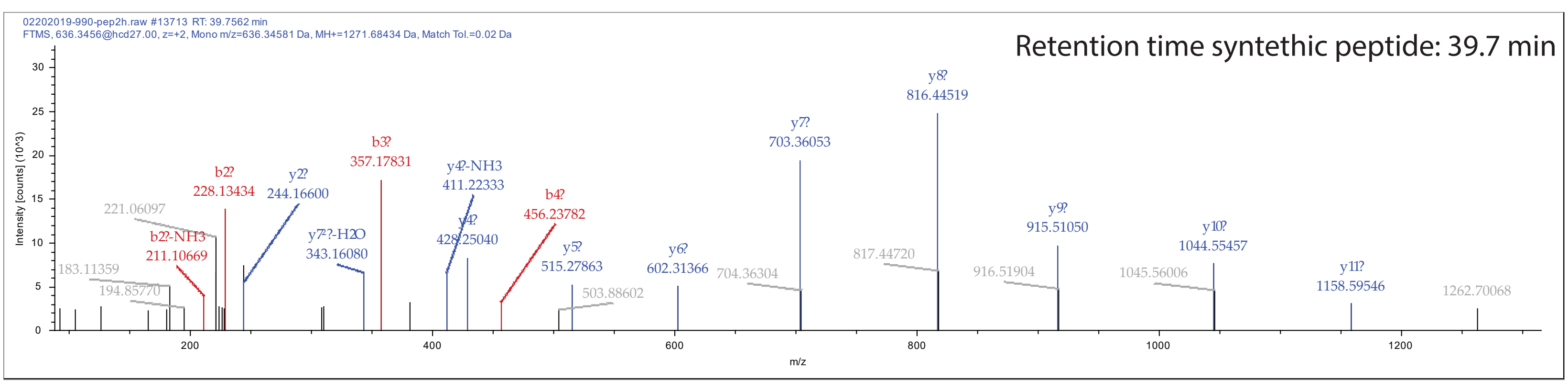


INIHLQILQK

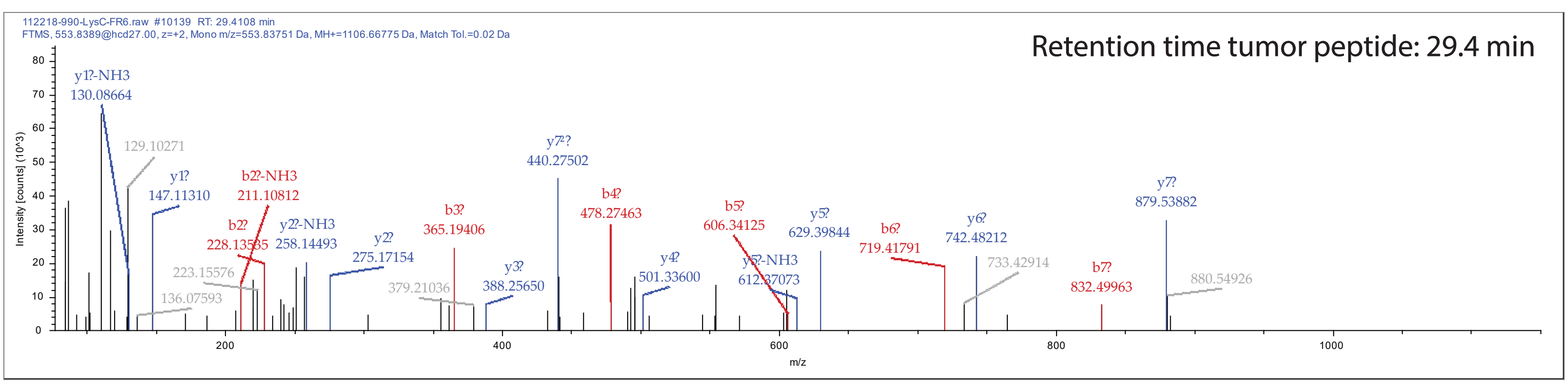


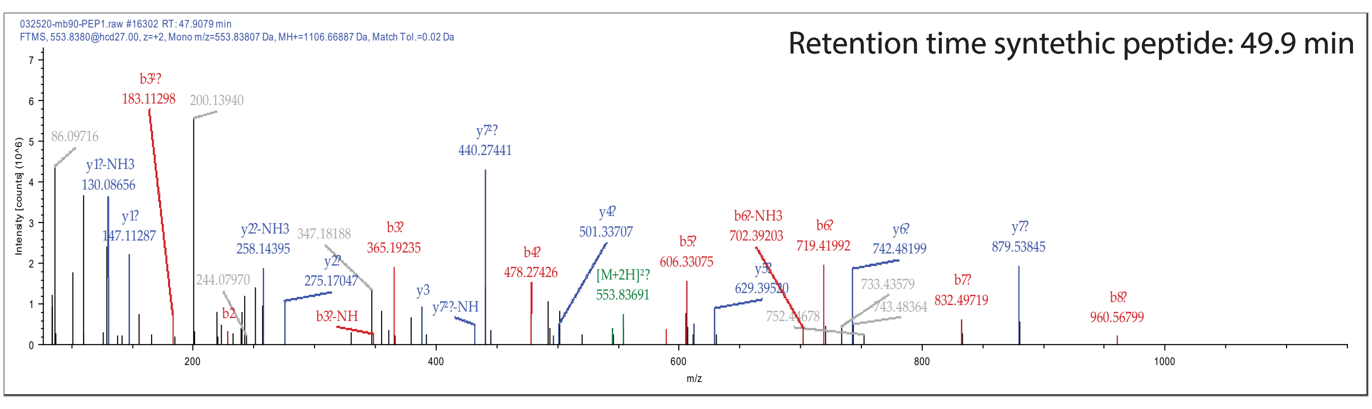


QGQGGNCDLRSSLGQGGK

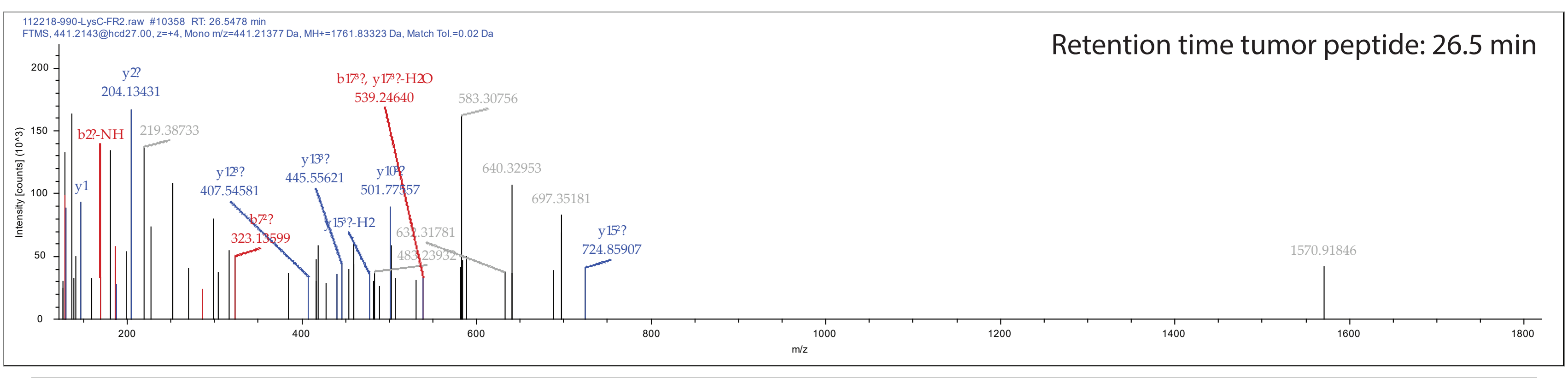


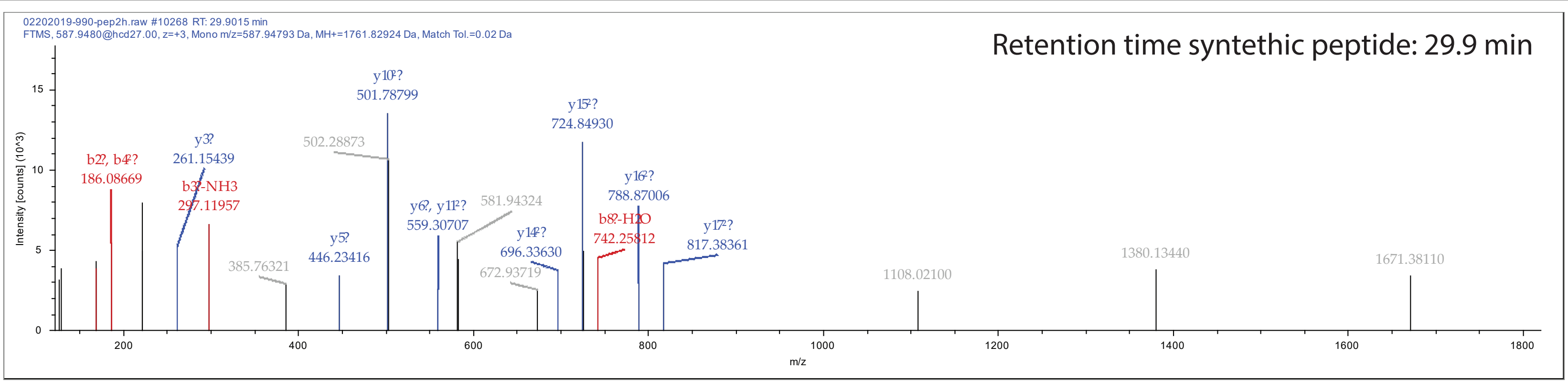


RLAGGFWCSPTRK

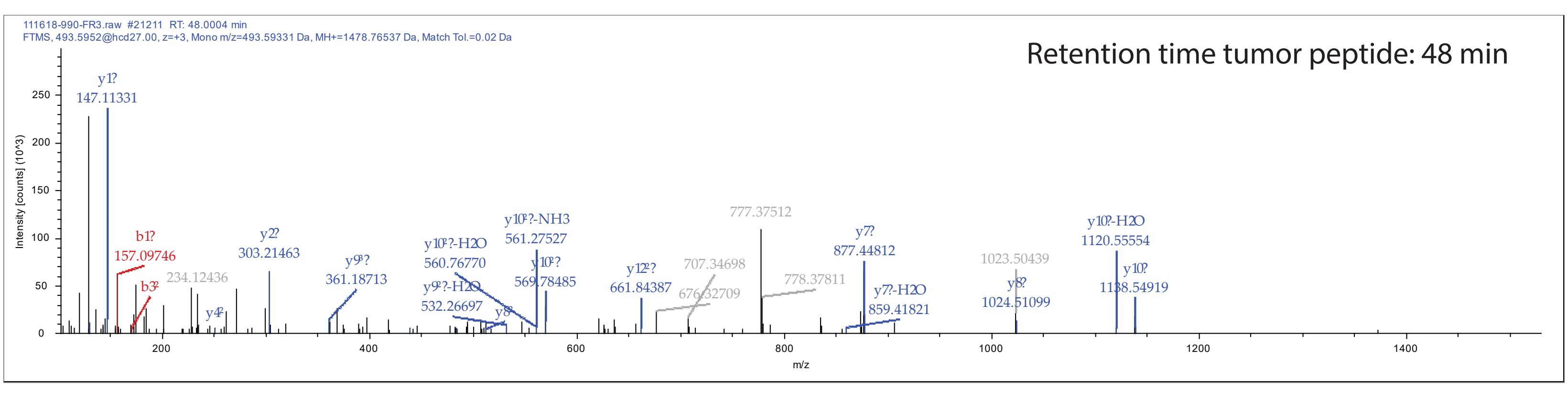


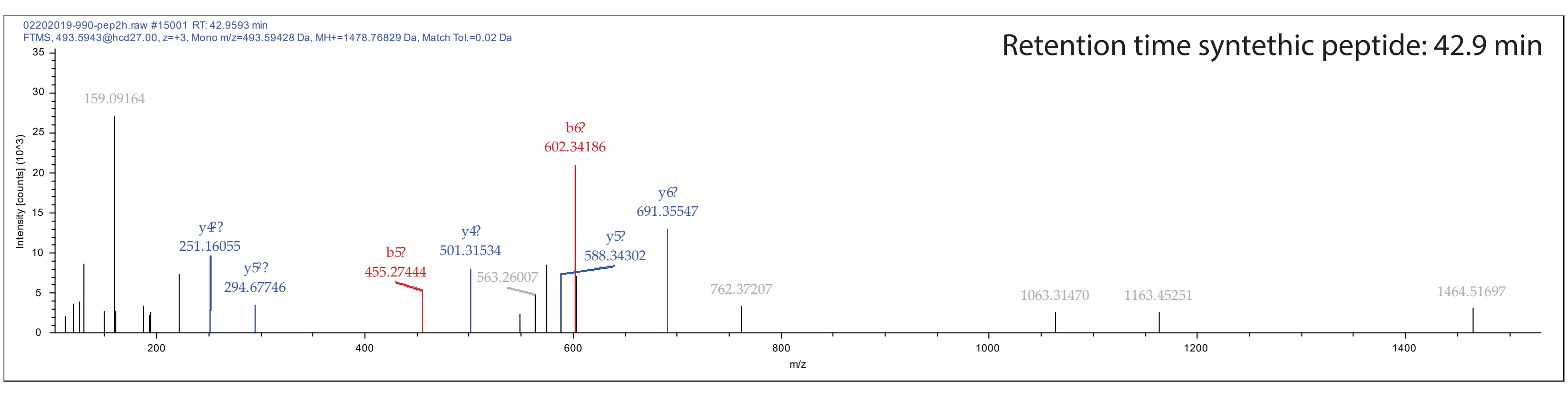


TTAGGESALAPSVFK

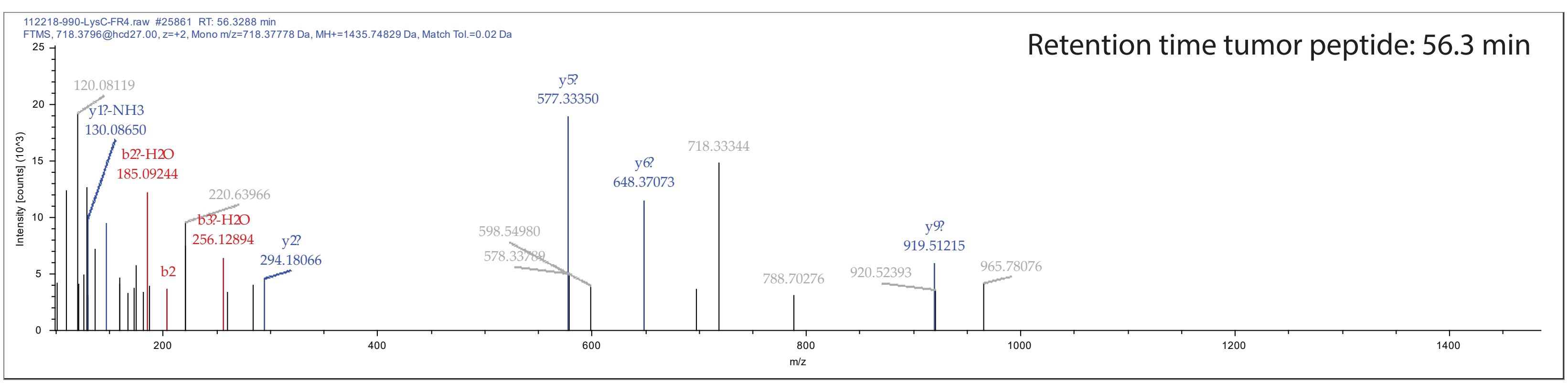


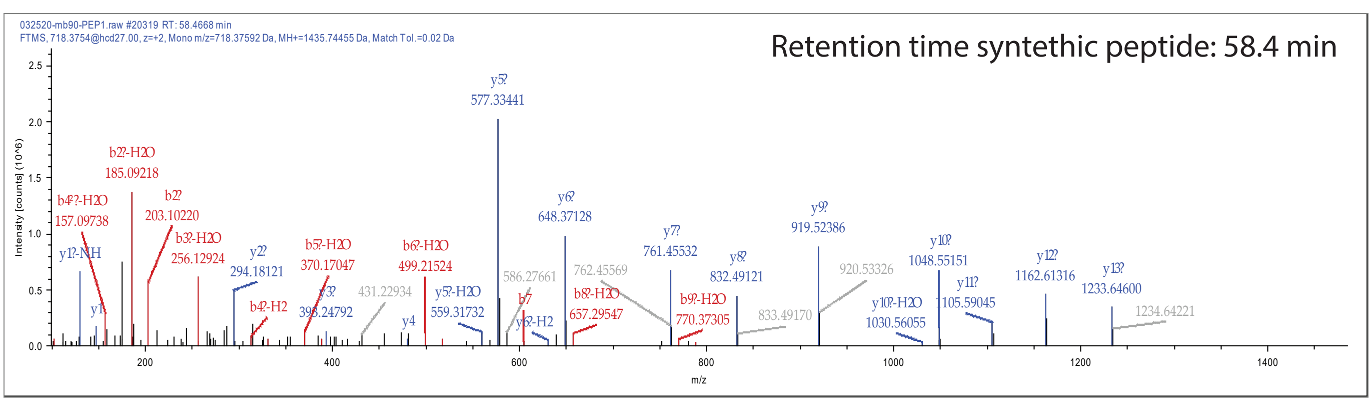


**Supplementary Fig. 6**

**Supplementary Fig. 6. PRM validation of peptides identified in the 7316-3778 primary tumor sample.** Retention time and spectra for the tumor tissue peptides and the synthetic peptides are shown.

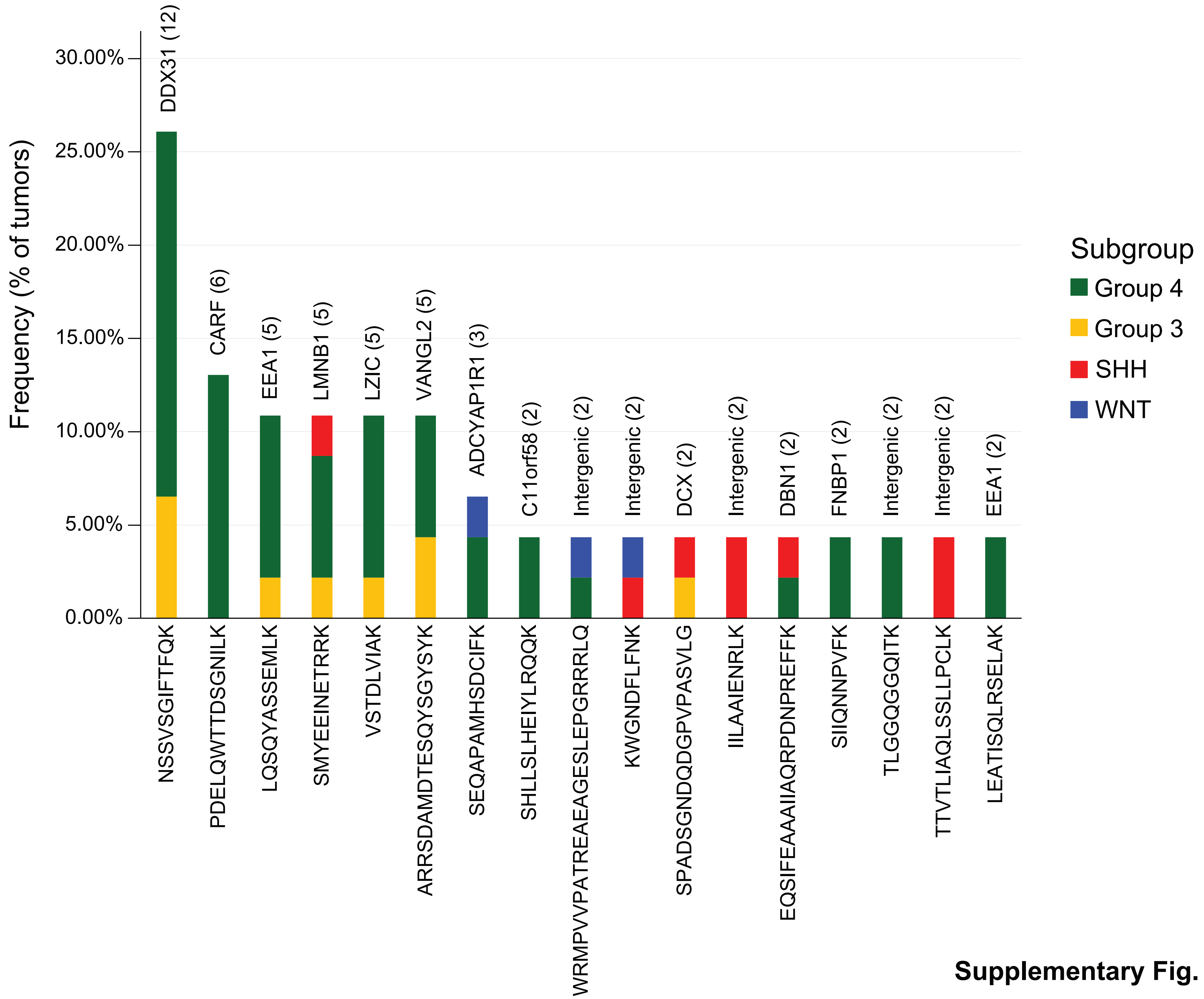


**Supplementary Fig. 7**

**Supplementary Fig. 7. Tumor specific peptide frequency.** Bar plot depicting the frequent novel peptides found in 46 medulloblastoma tumors. All shared peptides originate from novel junctions. The genes involved in the novel junctions are indicated in the figure. The numbers indicate the total number of tumors in which the novel peptides were found. Colors indicate the medulloblastoma subgroup; green, yellow, red, and blue for Group 4, Group 3, SHH and WNT subgroups respectively.

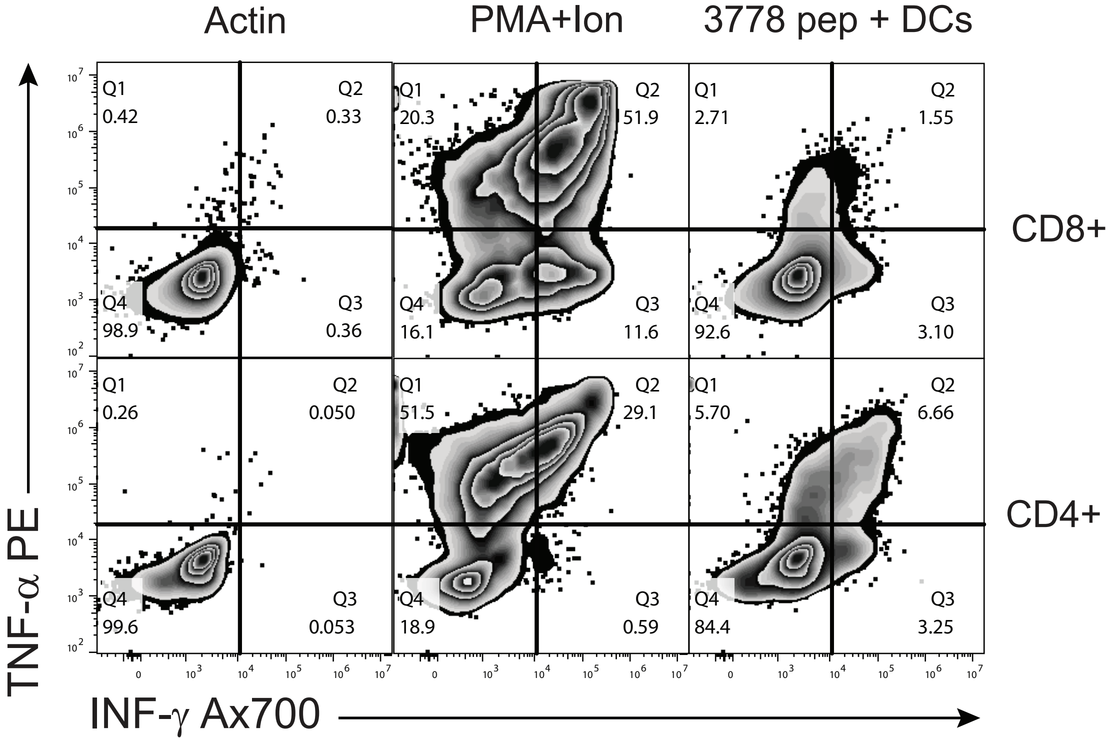


**Supplementary Fig. 8**

**Supplementary Fig. 8. T cell IFN-γ and TNF-α response from patient 7316-3778.** Autologous T cells were stimulated with 15 peptides identified in patient 7316-3778's tumor. Specific CD4+ T cell responses to pooled peptides were observed by intracellular flow cytometry. PMA+Iono: phorbol myristate acetate and ionomycin. Gating strategy in Supplementary Fig. 15.

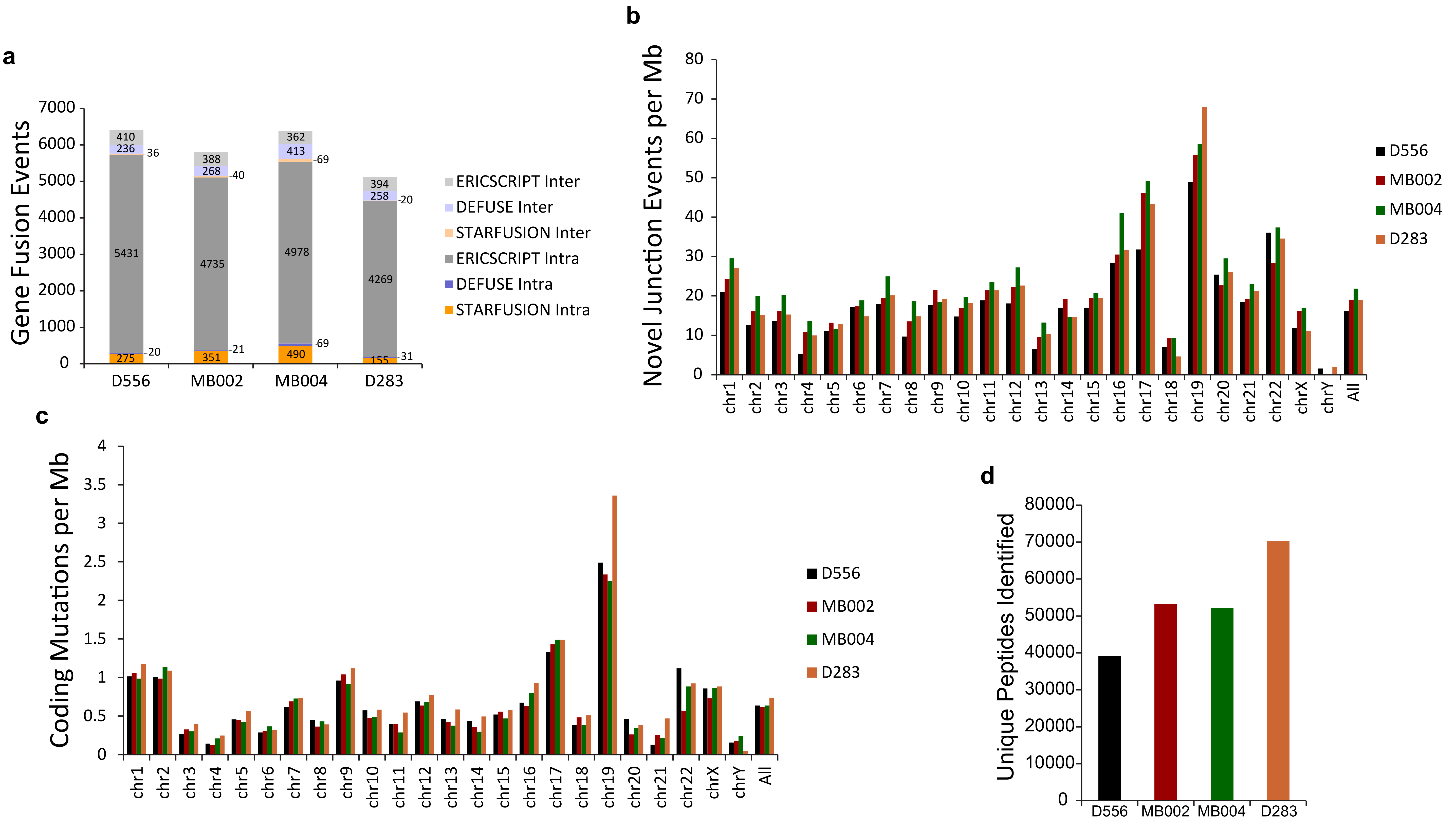


**Supplementary Fig. 9**

**Supplementary Fig. 9. Characterization of the genomic and proteomic events identified in Medulloblastoma cells lines. a** Aggregate barplot indicating the number of fusions detected in medulloblastoma cell lines. Inter and intra chromosomal fusions for each fusion caller are represented. **b** Barplot indicating the number of novel junctions per megabase identified in the medulloblastoma cell lines. **c** Barplot indicating the number of somatic coding mutations per megabase identified in the medulloblastoma cell lines. **d** Barplot indicating the total number of unique peptides identified by LC-MS/MS in medulloblastoma cell lines.

Peptide Validation MB002

DLSLVTER

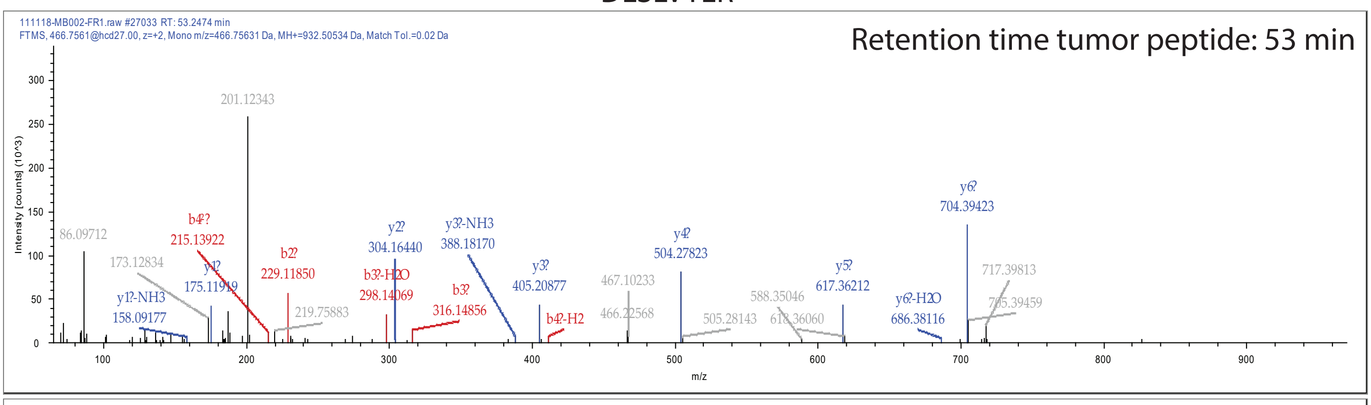


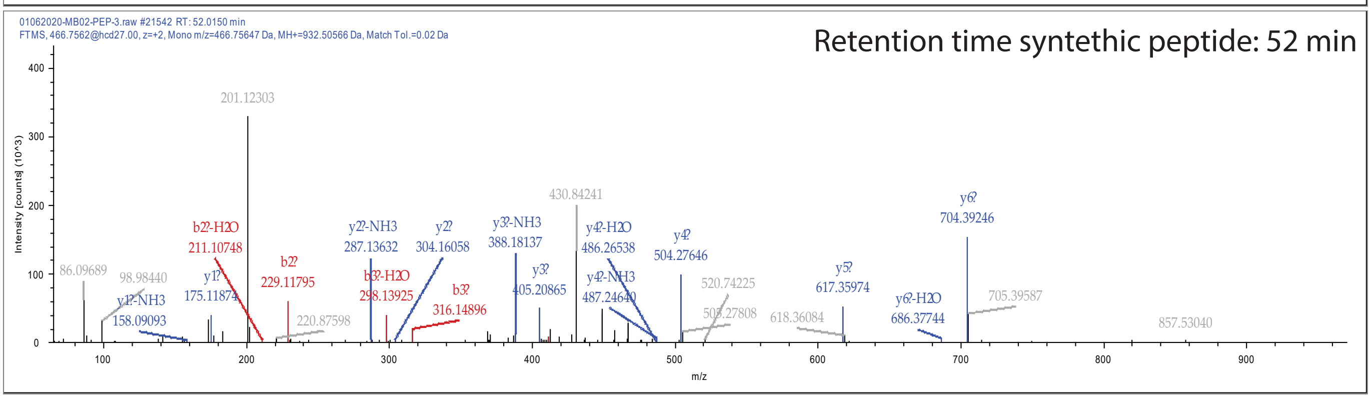


KEYNADYDLSAR

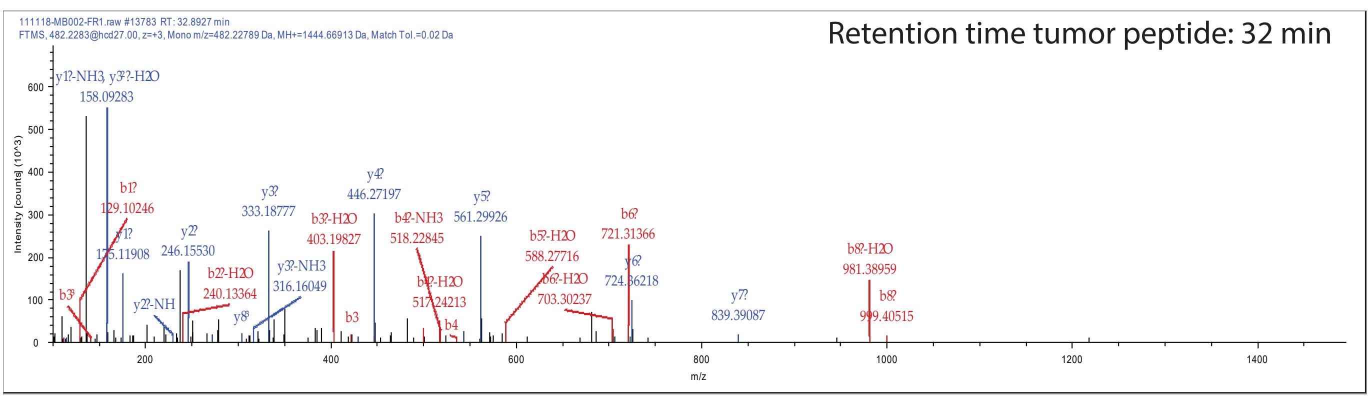


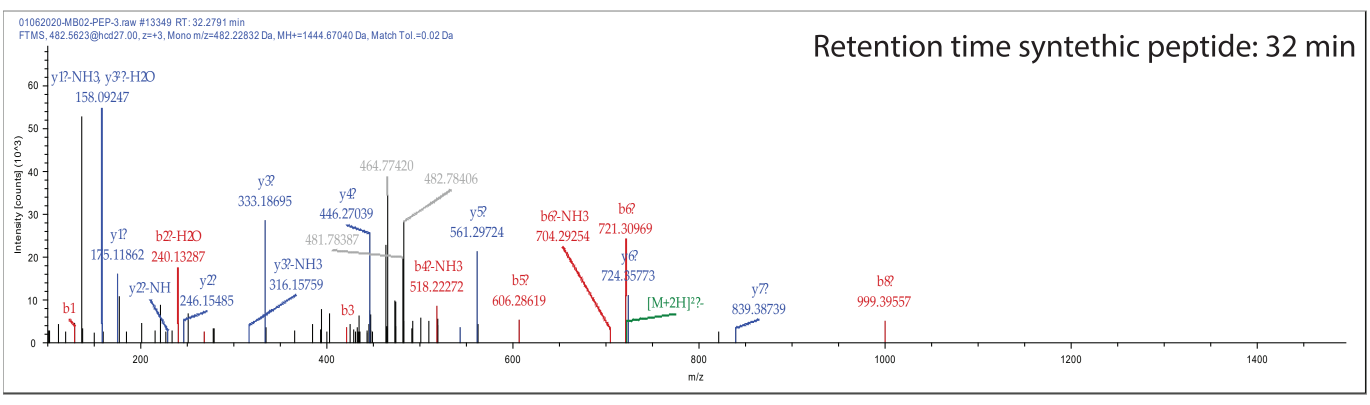


LDLAGNLPGSK

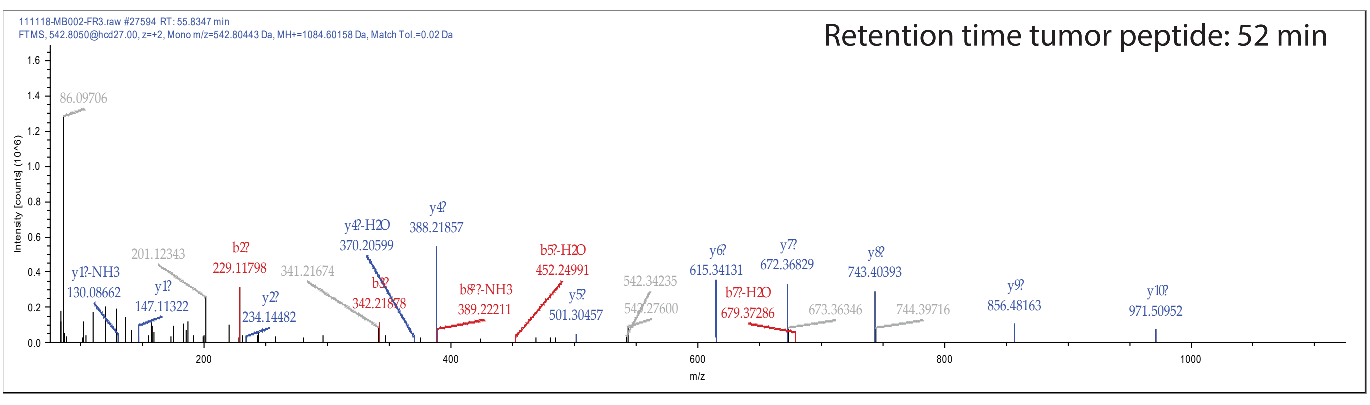


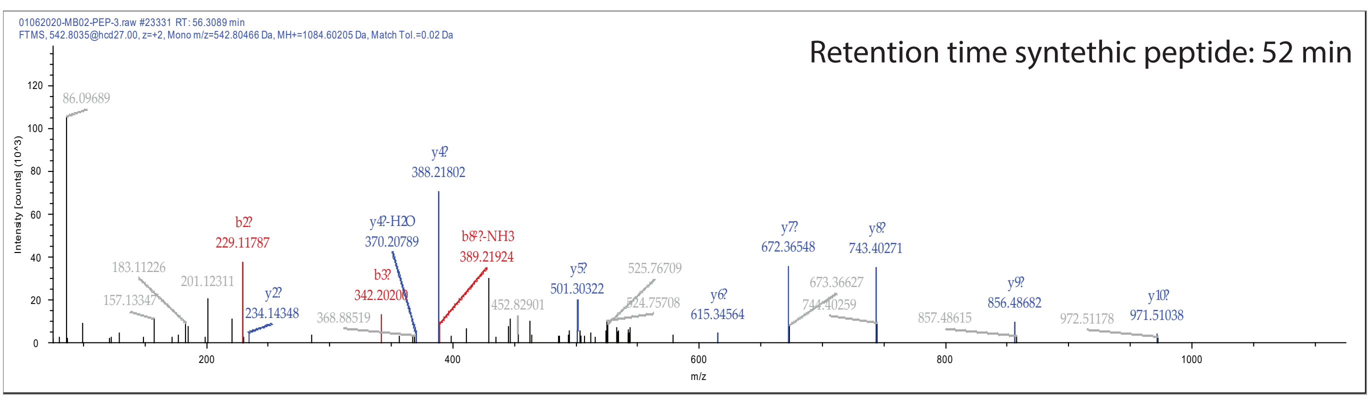


PAFDLFENR

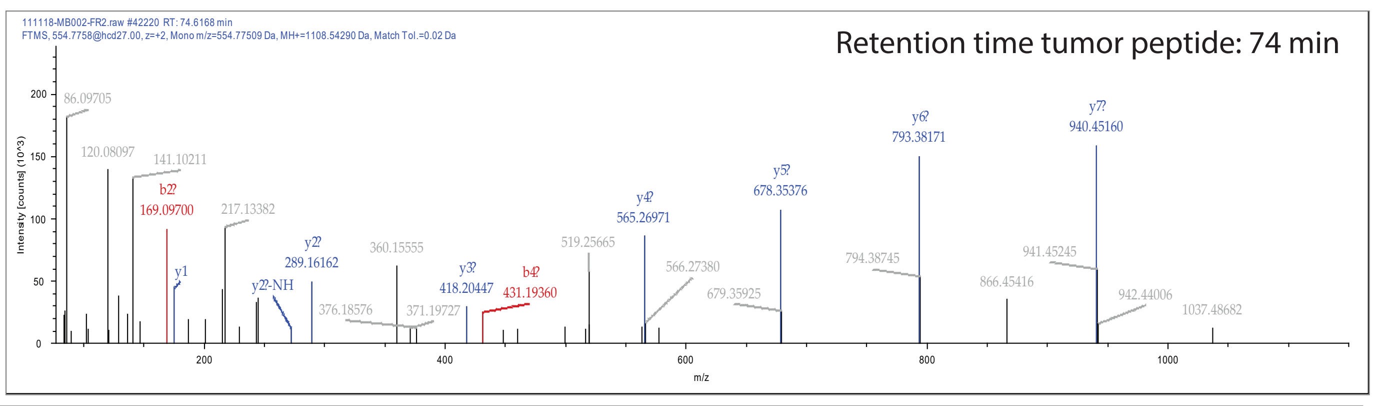


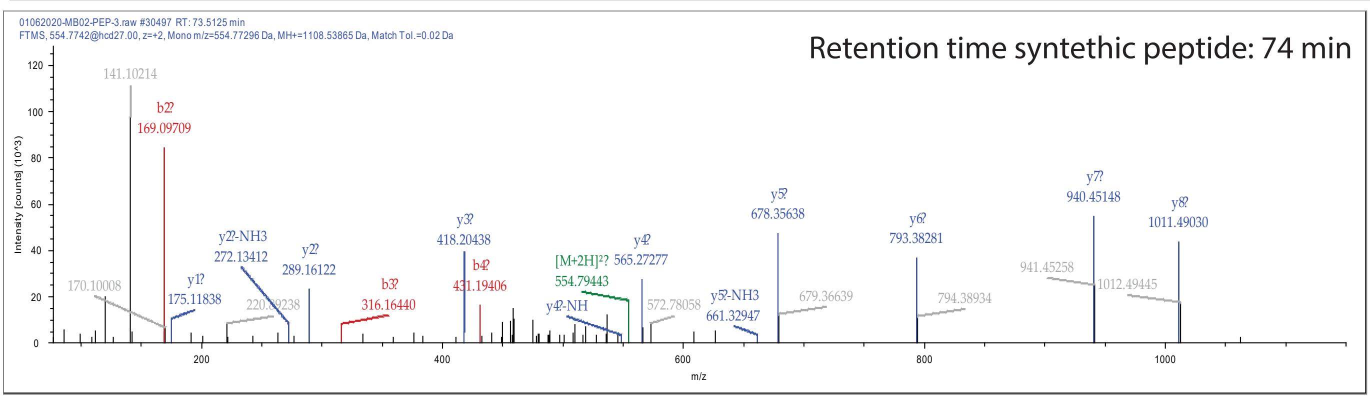


RFDSELSQAR

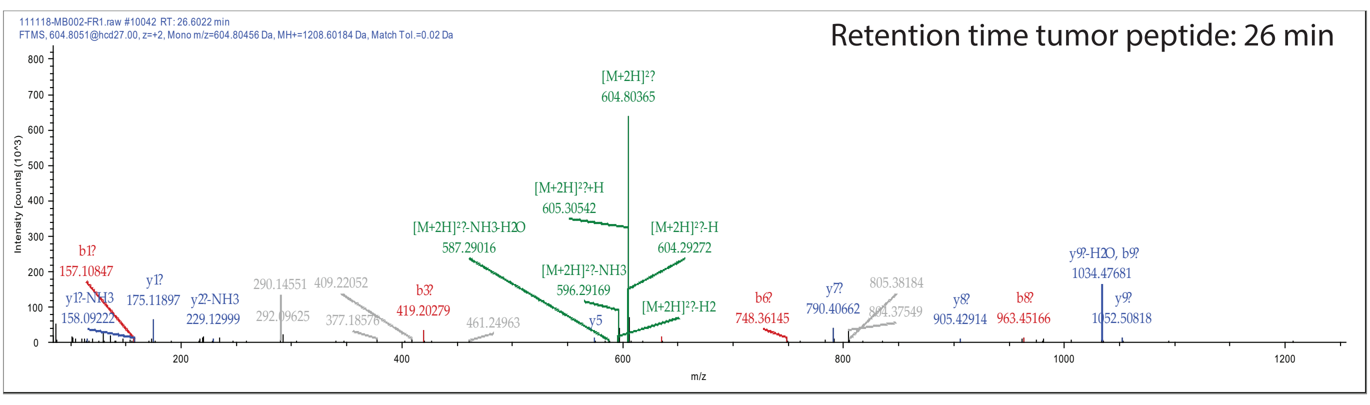


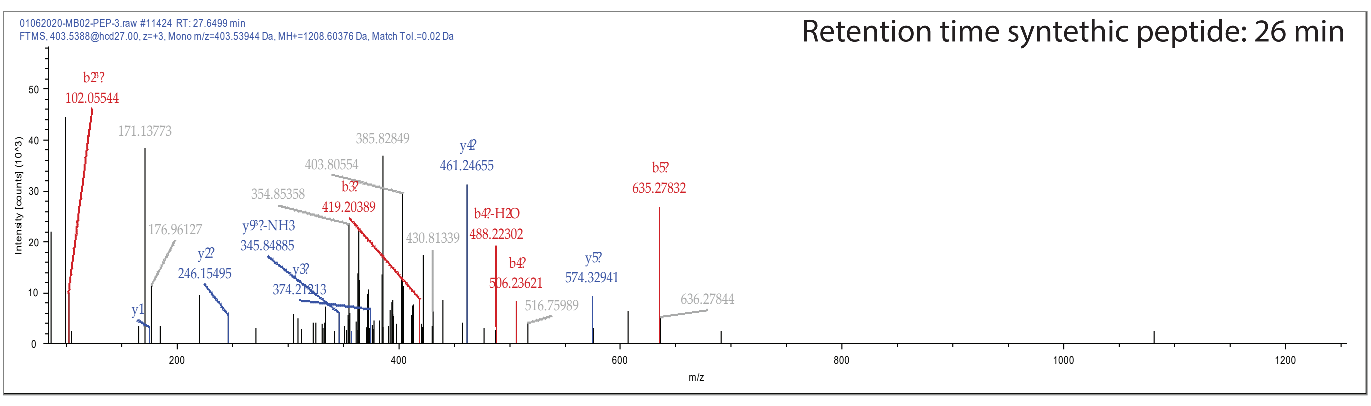


VLLLSIQNPLYPITVPTR

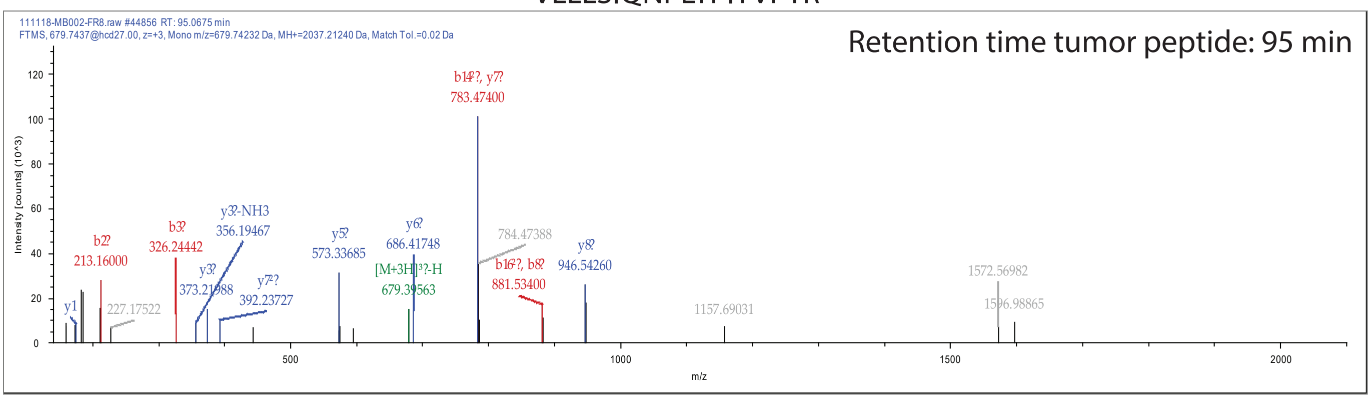


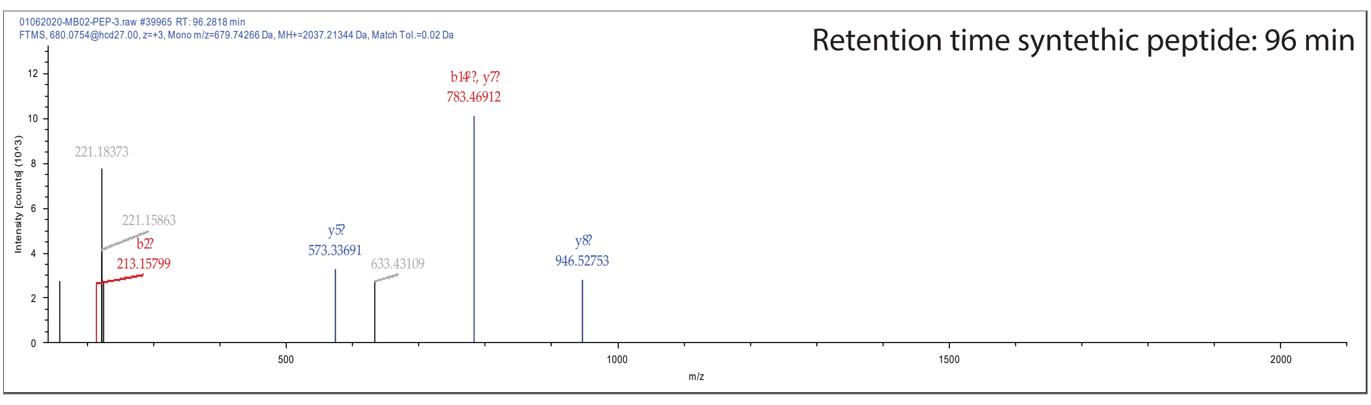


**Supplementary Fig. 10**

**Supplementary Fig. 10. PRM validation of peptides identified in the MB002 cell line.** Retention time and spectra for the cell line peptides and the synthetic peptides are shown.

Peptide Validation D556

DLTQTASSTARGEK

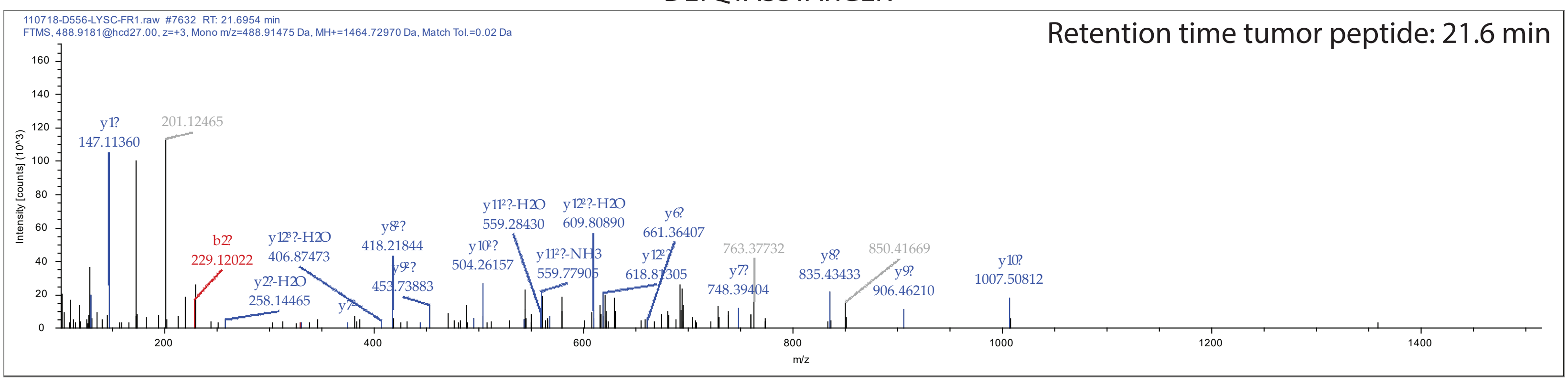


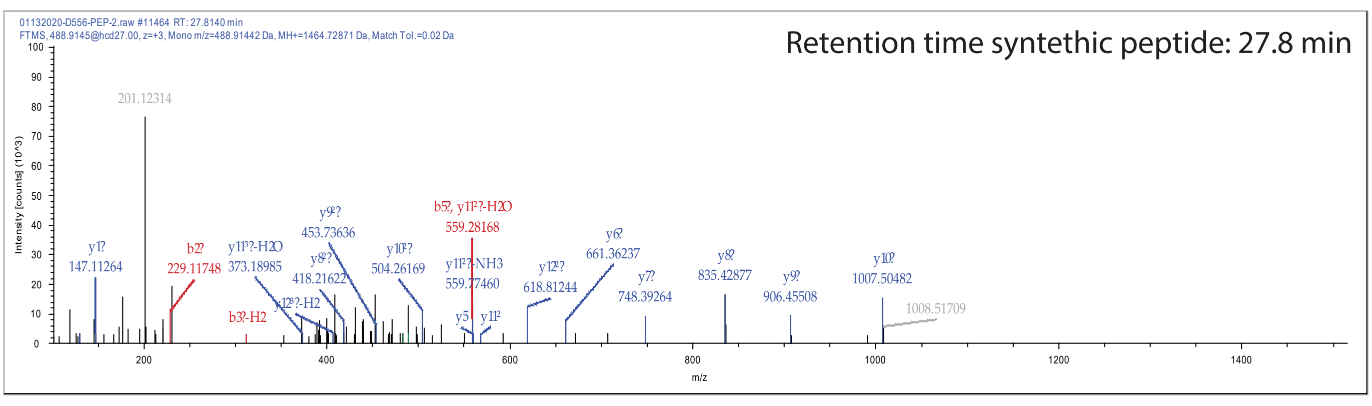


ITGISFIPR

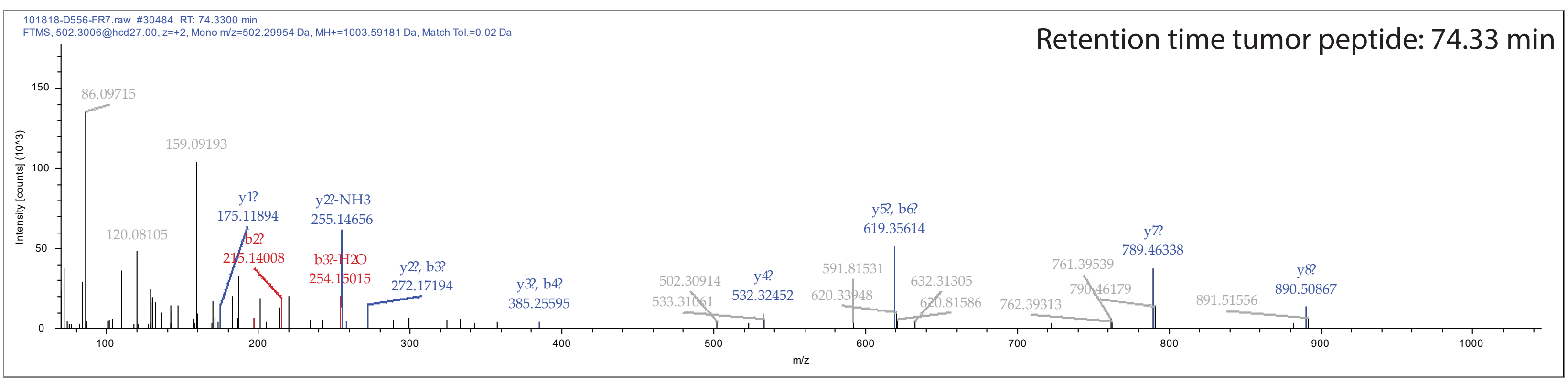


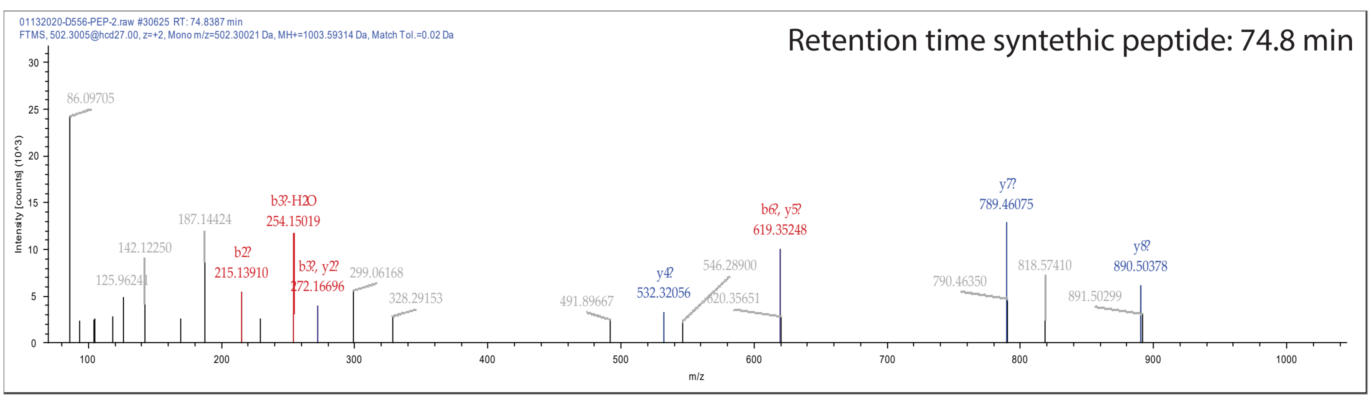


KEYNADYDLSAR

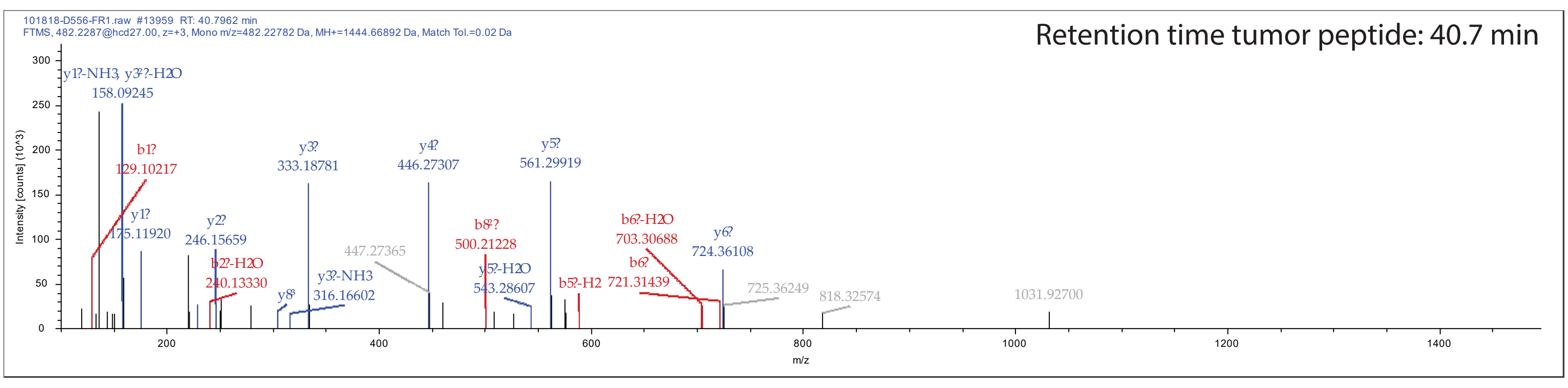


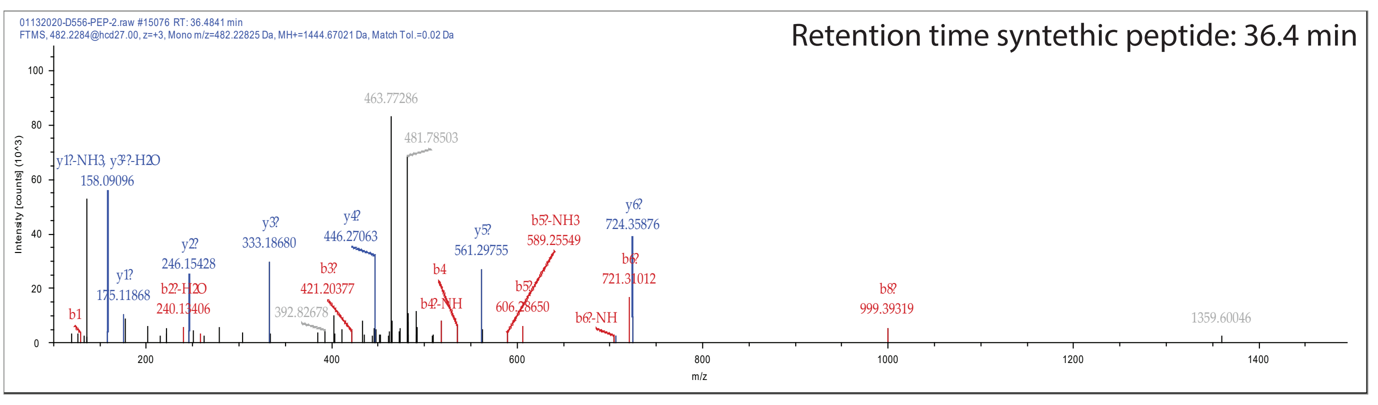


LSLEYRPVIDK

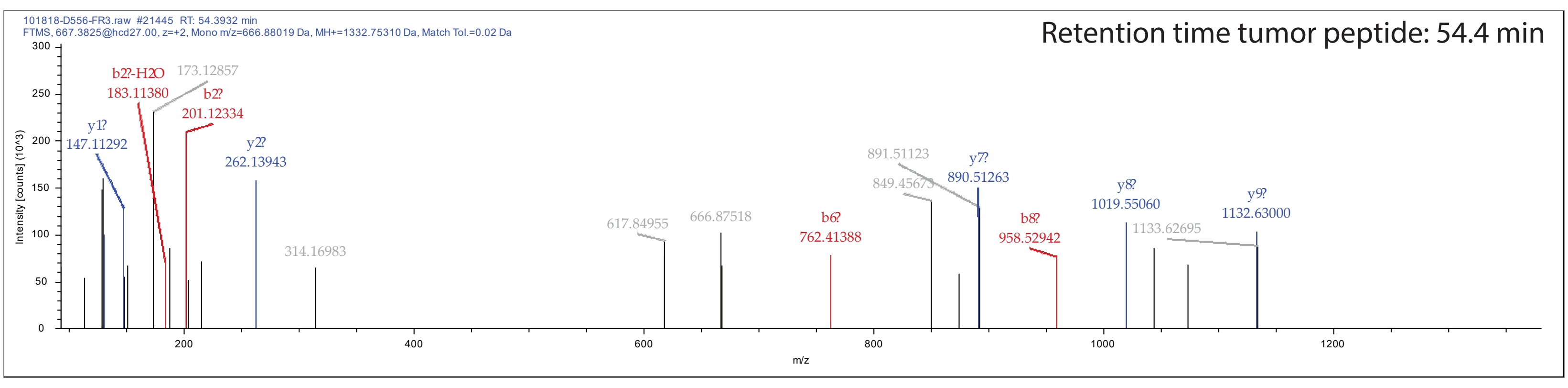


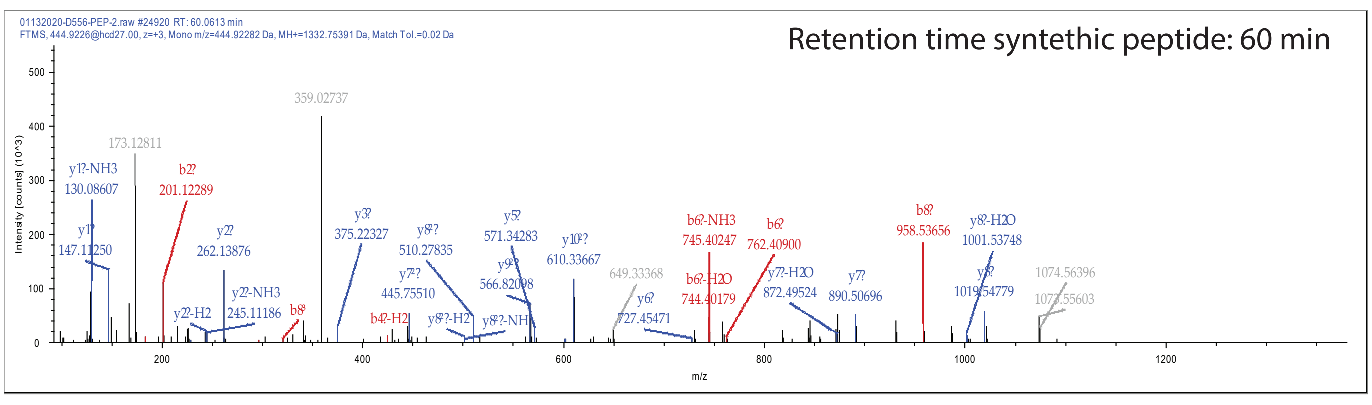


QSLLLNGIGVR

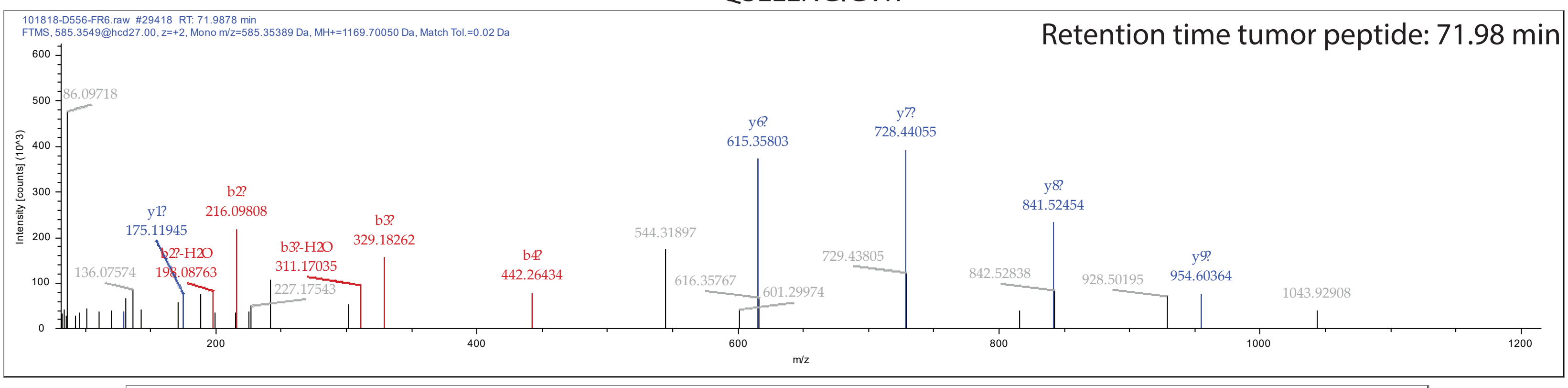


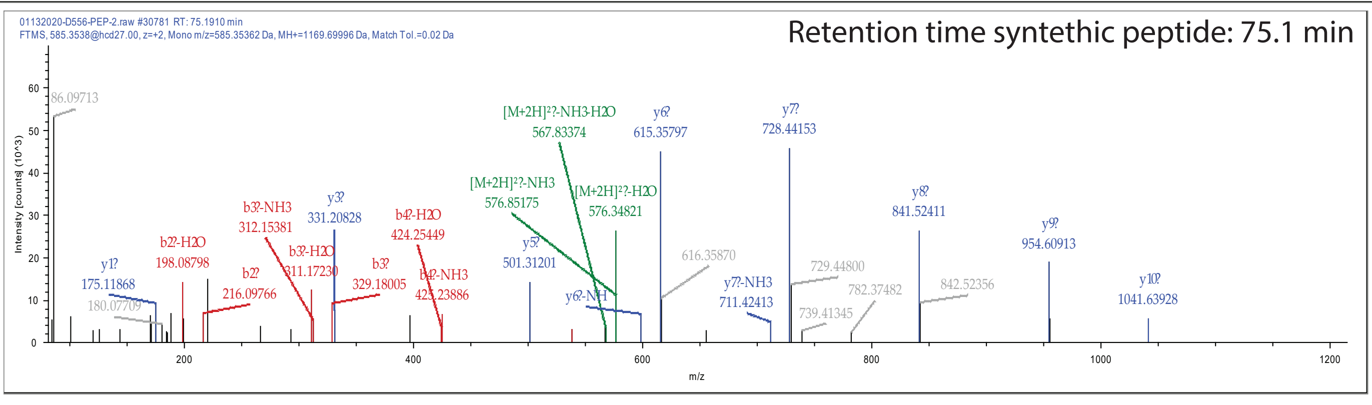


VDIDTPDIDIHGPDAK

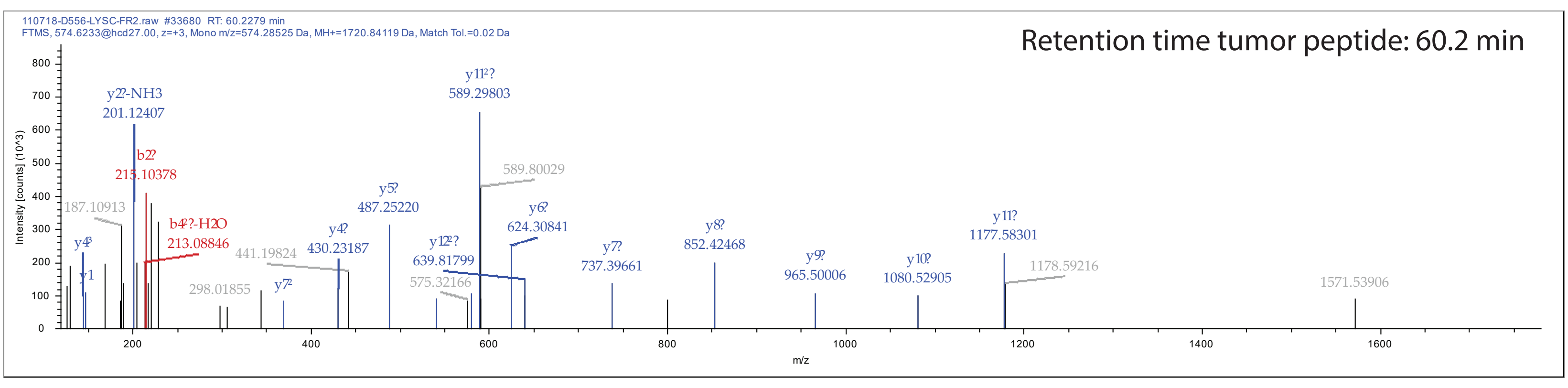


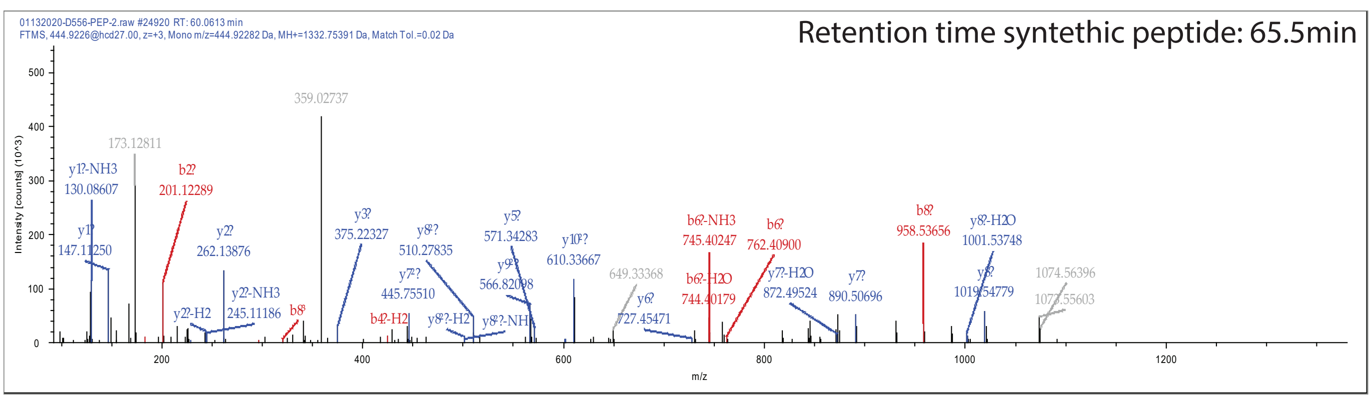


**Supplementary Fig. 11**

**Supplementary Fig. 11. PRM validation of peptides identified in the D556 cell line.** Retention time and spectra for the cell line peptides and the synthetic peptides are shown.

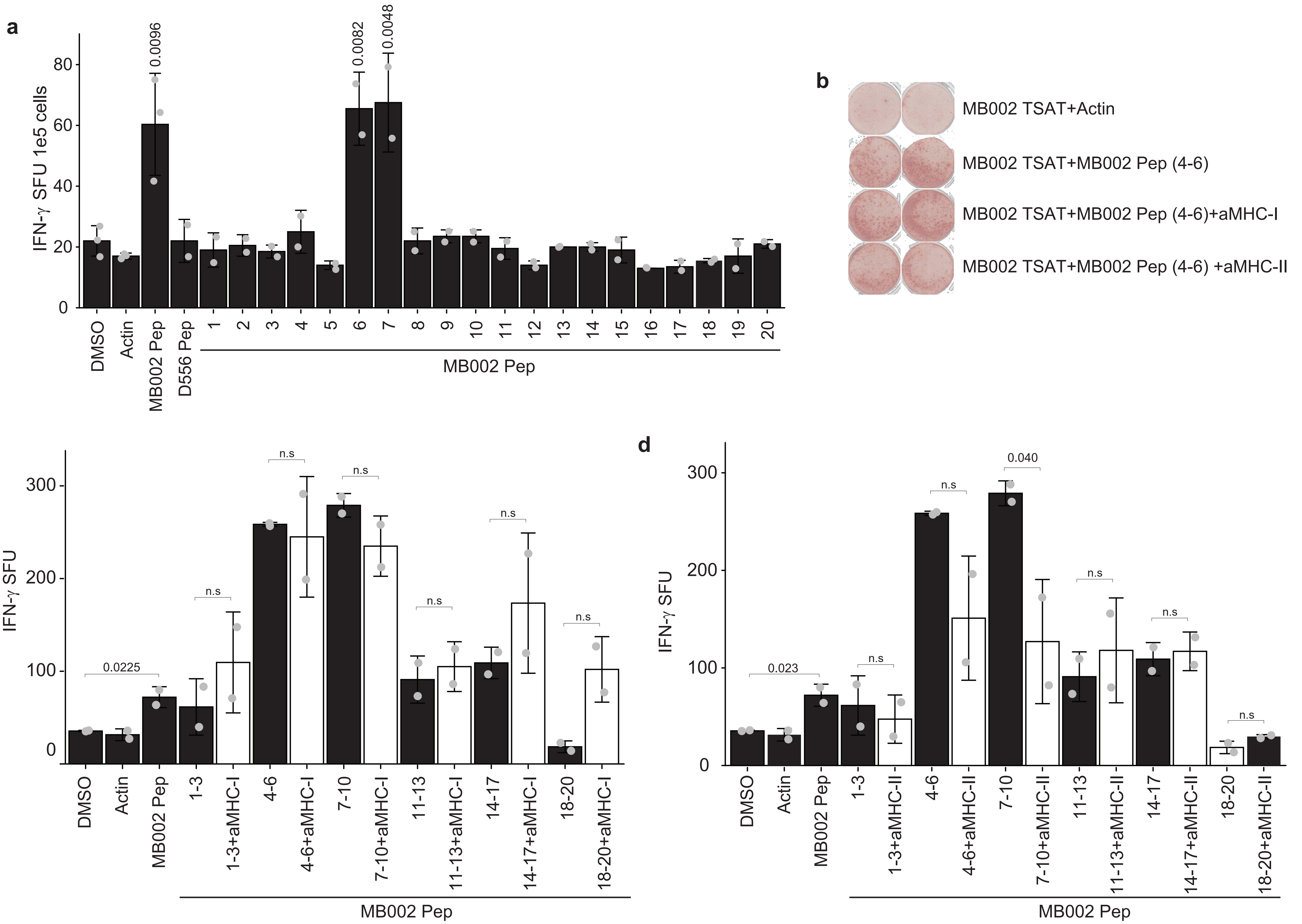


**Supplementary Fig. 12**

**Supplementary Fig. 12. MB002 cell line specific peptides induce peptide-specific T cell IFN-γ responses *in vitro*.** Dendritic cells were loaded with MB002-specific peptides and co-cultured with non-adherent PBMCs. After 3 stimulations, peptide-specific T cell responses were analyzed by anti-IFN-γ ELISpot. **a** Summary data of IFN-γ response to pooled MB002 peptides and peptides 6 & 7 in the healthy donor 3. One-sided t-test was used to calculate the p-values, n=2. p-values are indicated the figure. **b** Representative ELISpot membranes showing positive response to MB002 peptide pools and attenuated response in the presence of MHC-II blocking antibodies. **c, d** Summary data of responses to peptide sub-pools in the presence of MHC Class I and II blocking antibodies in healthy donor 1. One-sided t-test was used to calculate the p-values, n=2. p-values are indicated the figure. SFU: spot-forming units; 1 SFU = 1 T cell secreting IFN-γ; TSAT: Tumor-specific antigen T cell; a-MHC-I, a-MHC-II: anti-MHC Class I and II blocking antibodies; MB002 pep: pooled MB002 peptides; D556 pep: pooled D556 peptides; DMSO: dimethyl sulfoxide (peptide solvent; unstimulated control); actin: peptide specificity control. Error bars represent one standard deviation (SD).

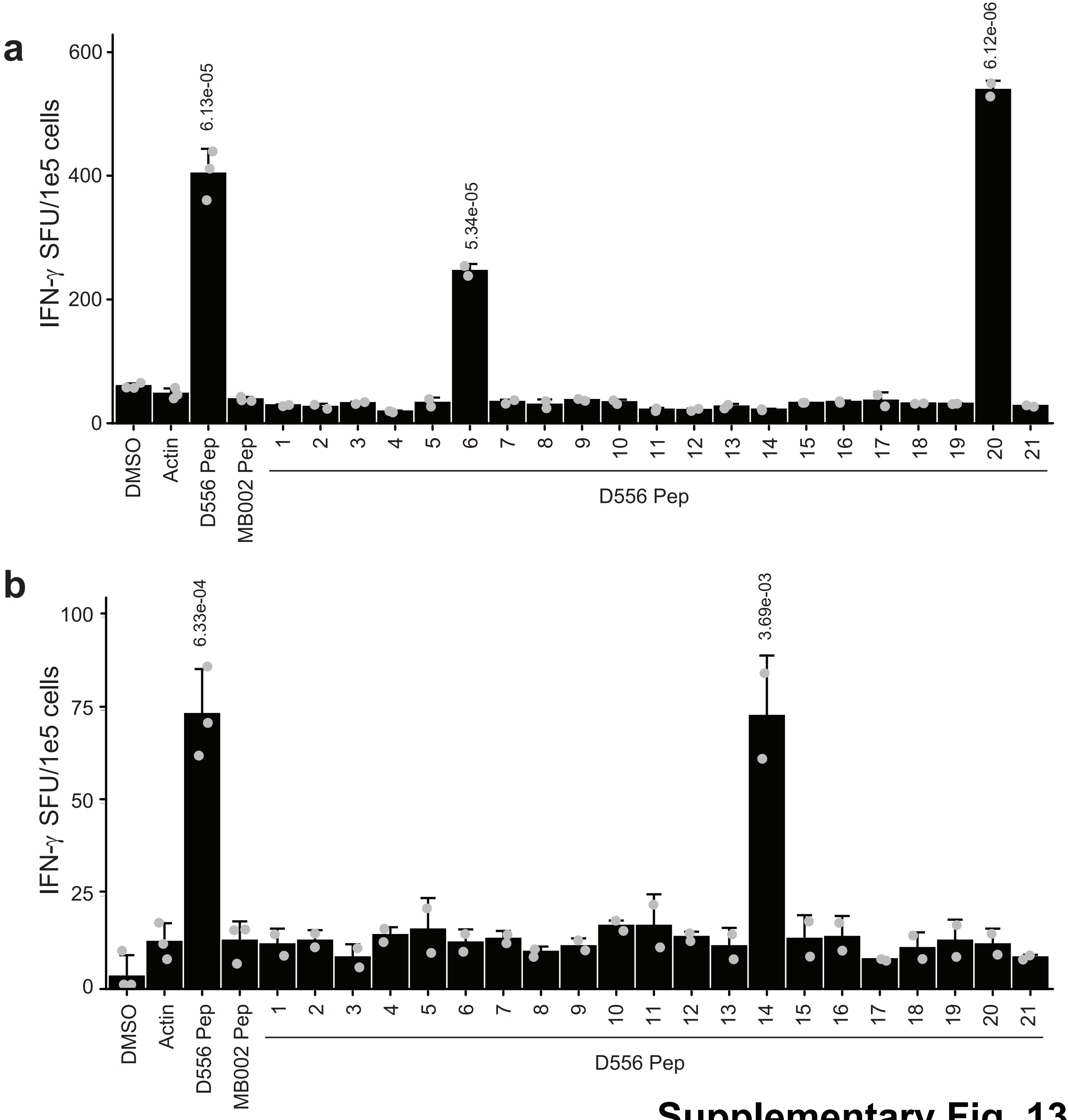


**Supplementary Fig. 13**

**Supplementary Fig. 13. D556 cell line-specific peptides induce specific T cell IFN-γ responses *in vitro***. Dendritic cells were loaded with MBD556-specific peptides and co-cultured with non-adherent PBMCs. After 3 stimulations, peptide-specific T cell responses were analyzed by anti-IFN-γ ELISpot. **a, b** Summary data of anti-IFN-γ response by MB D556-primed T cells in 2 healthy donors (4 and 5), showing response to peptides 6 and 20 in donor 1 and peptide 14 in donor 2. One-sided t-test was used to calculate the p-values, n=2. p-values are indicated the figure. SFU: spot-forming units; 1 SFU = 1 T cell secreting IFN-γ; MB002 pep: pooled MB002 peptides; D556 pep: pooled D556 peptides; DMSO: dimethyl sulfoxide (peptide solvent; unstimulated control); actin: peptide specificity control. Error bars represent one standard deviation (SD).

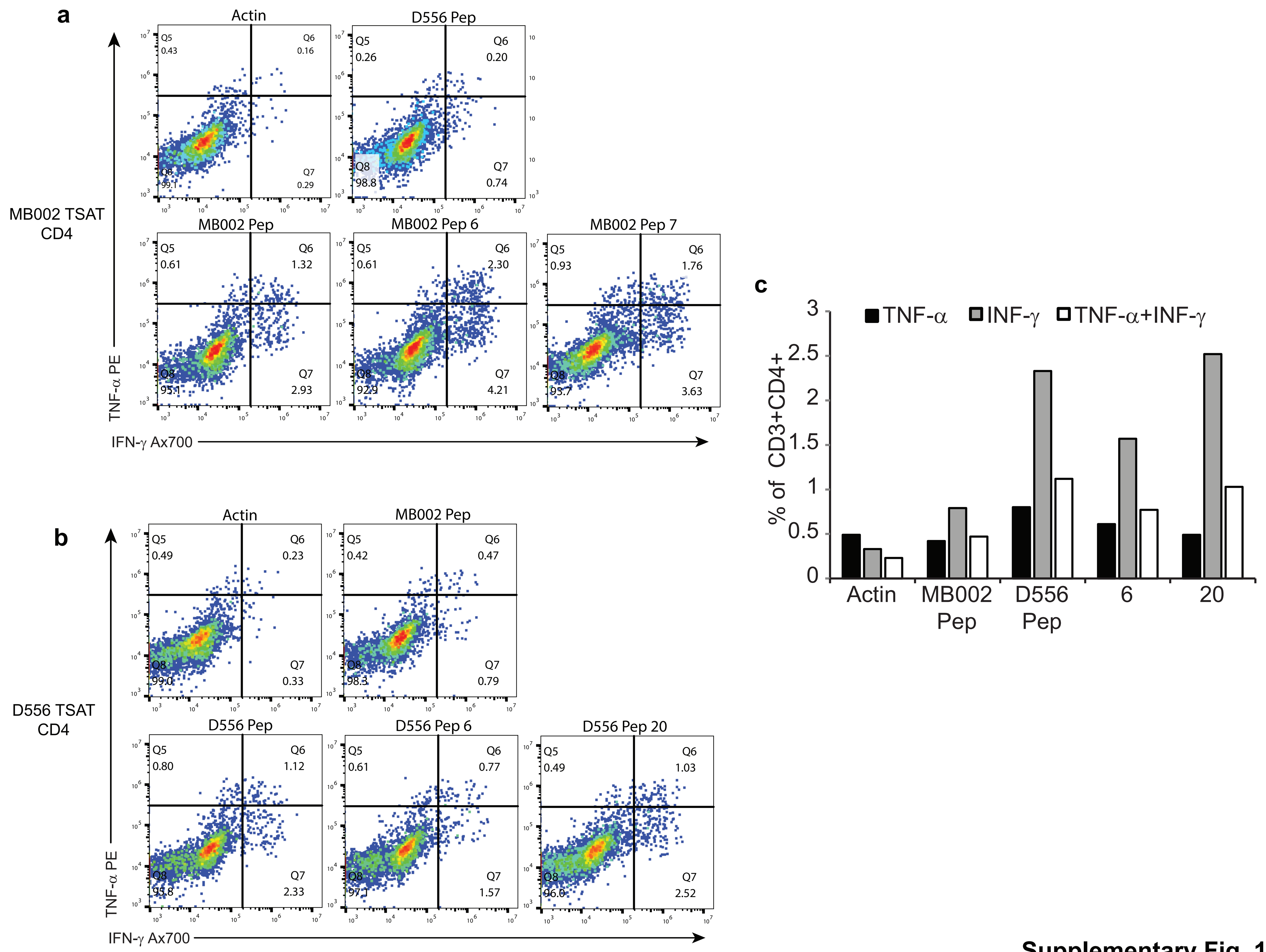


**Supplementary Fig. 14**

**Supplementary Fig. 14. Peptide-specific polyfunctional CD4+ MB002 and D556 TSAT can be identified in TSAT populations**. To assess cytokine function, TSAT were incubated in the presence of pooled TSA peptides. **a** Representative dot plots from healthy donor 2 showing specific IFN-γ and TNF-α responses to pooled and individual MB002 peptides by CD4+ TSAT. **b** Representative dot plots from healthy donor 4 showing specific IFN-γ and TNF-α responses to pooled and individual MB D556 peptides by CD4+ TSAT. **c** Summary data of Flow cytometry CD4+ response to D556 peptides 6 and 20. MB002 pep: pooled MB002 peptides; D556 pep: pooled D556 peptides. Gating strategy in Supplementary Fig. 15.

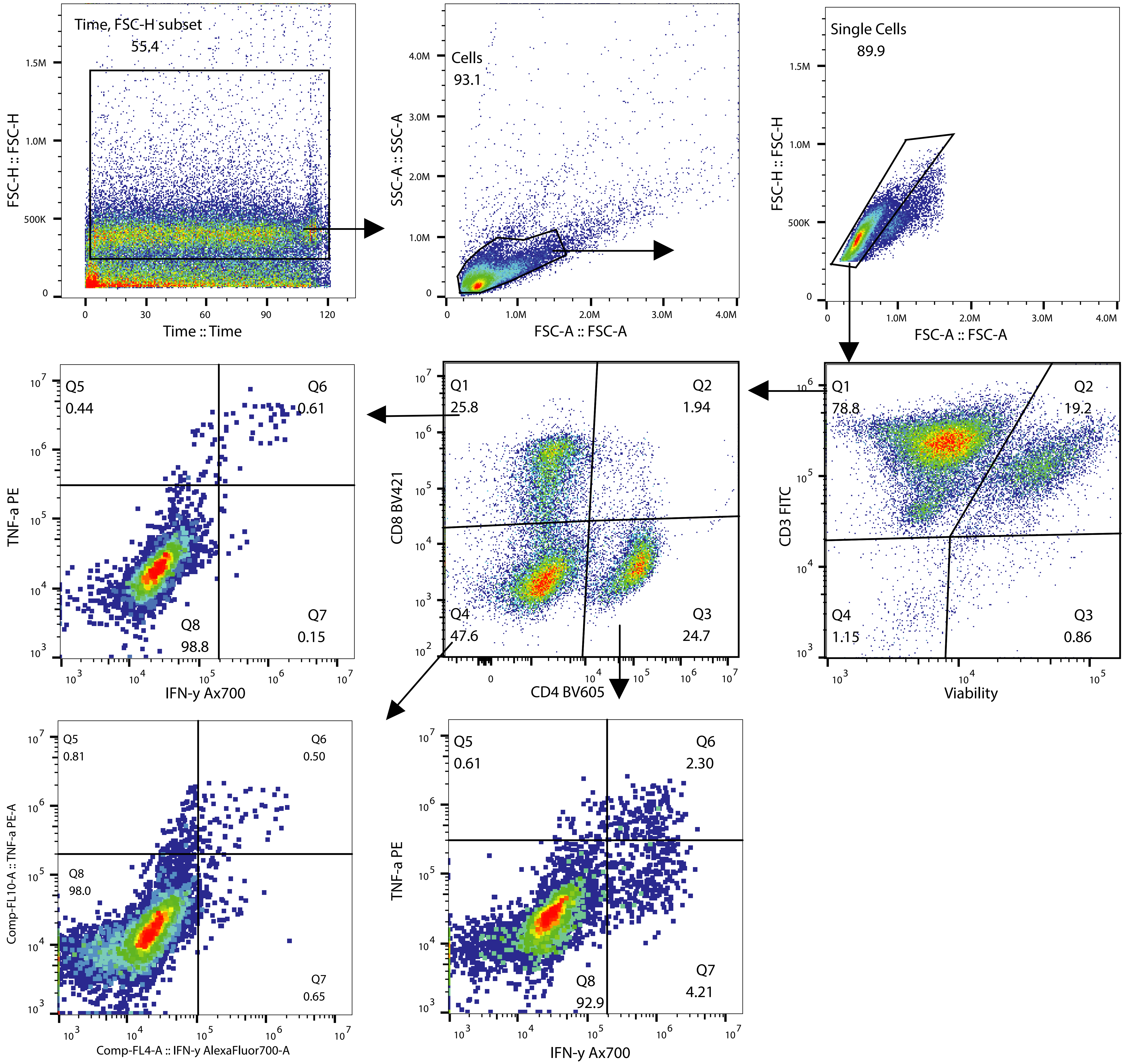


**Supplementary Fig. 15**

**Supplementary Fig. 15. Gating strategy for intracellular cytokine expression**. Dendritic cells were loaded with MB-specific peptides and co-cultured with non-adherent PBMCs. After 3 stimulations, peptide-specific T cell populations were characterized by Flow Cytometry. Cells were labeled with FcR block, dead cell exclusion dye, surface and intracellular antibodies as in methods section (see supplementary dataset 10 for antibodies and panels). Cells were acquired on a Beckman Coulter CytoFlex S using CytExpert version 2.2.0.97 software. Data were analyzed on Flow Jo version 10.5. The gating tree for detecting intracellular cytokine expression in CD8+, CD4+ and CD8-CD4- cells is shown.

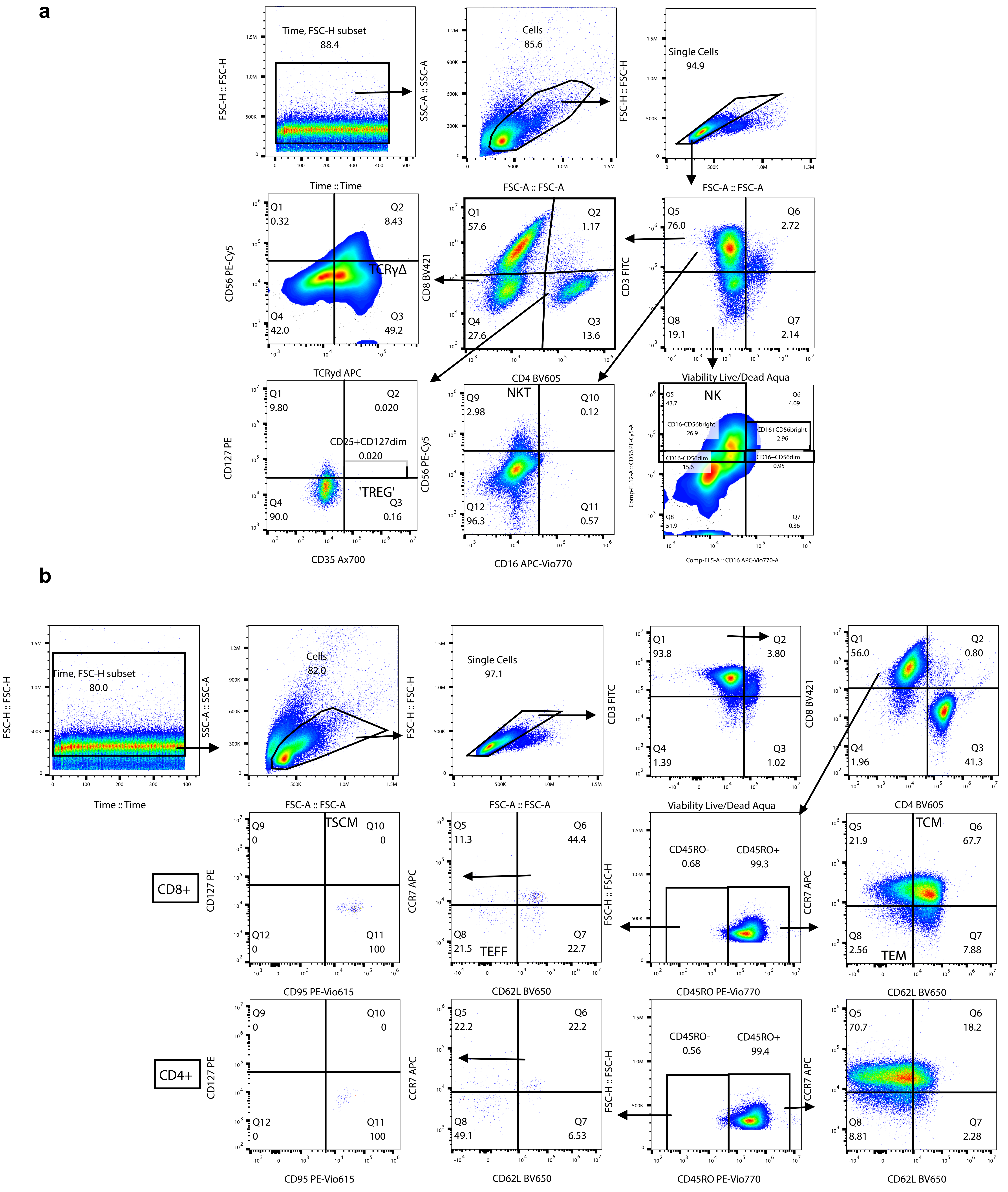


**Supplementary Fig. 16**

**Supplementary Fig. 16. Gating strategy for expanded T cell phenotype (a) and differentiation status (b)**. Dendritic cells were loaded with MB-specific peptides and co-cultured with non-adherent PBMCs. After 3 stimulations, peptide-specific T cell populations were characterized by Flow Cytometry. Cells were labeled with FcR block, dead cell exclusion dye and antibodies per the methods (see supplementary dataset 10 for antibodies and panels). Cells were acquired on a Beckman Coulter CytoFlex S using CytExpert version 2.2.0.97 software. Data were analysed on Flow Jo version 10.5. **a** The gating tree for detecting CD3+CD4+CD8+TCRγδ+, $T_{REGs}$, NK and NKT cells. **b** The gating tree for detecting T central memory (TCM), T effector memory (TEM), T effector (TEFF) & T stem cell memory (TSCM) cells.

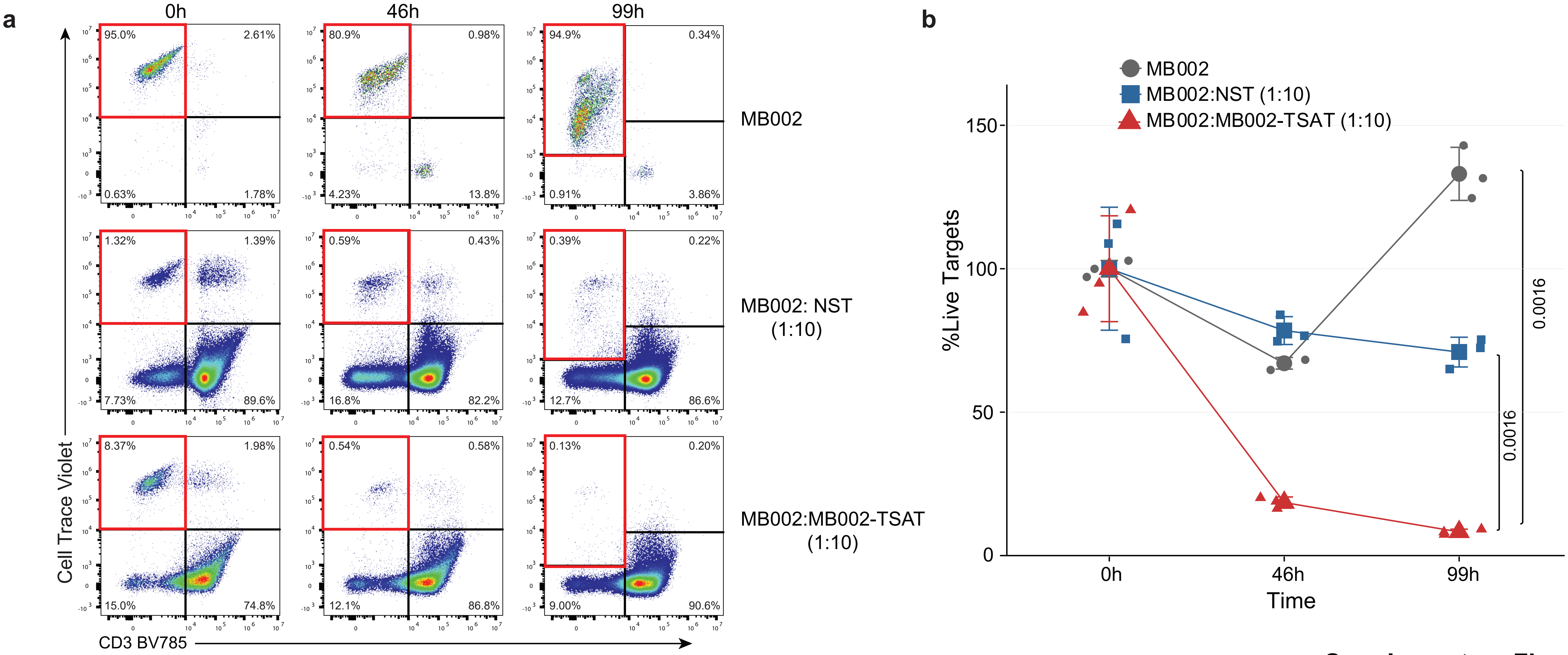


**Supplementary Fig. 17**

**Supplementary Fig. 17. MB002 cells are killed by TSA T and by non-specific T cells.** To assess cytotoxic function cryopreserved TSA T were thawed and rested overnight in medium containing IL-2 (100 U/mL). The following morning TSA T were plated 10:1 with target cells. At the indicated time points, co-cultures were stained as described in Materials & Methods. **a** Representative dot plots from the healthy donor 2 showing proliferation of MB002 targets in the absence of TSA T (top row), moderate disappearance (lysis) of tumor targets in the presence of non-specifically activated T cells (PHA blasts; middle row) and robust lysis of tumor targets in the presence of TSA T (bottom row). Lysis was determined based on the disappearance of targets from quadrant 1 (red border). **b** Summary data of A. NST: non-specific T cells (PHA blasts). One-way ANOVA p-values are shown. Values at each time point were normalized to 0 hours (100%). NST: non-specific T cells (PHA blasts). Gating strategy for expanded TSA T tumor cell cytotoxicity assays (Supplementary Fig. 18). Error bars represent one standard deviation (SD).

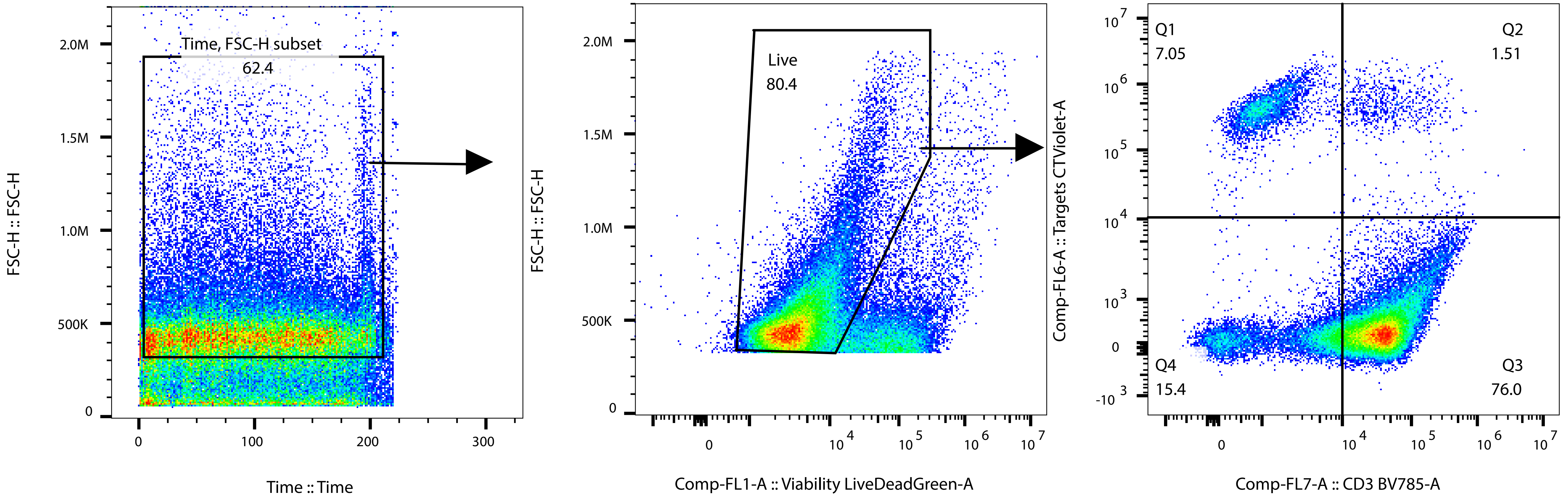


**Supplementary Fig. 18**

**Supplementary Fig. 18. Gating strategy for expanded TSA T tumor cell cytotoxicity assays.** To assess cytotoxic function, cryopreserved TSA T were thawed and rested overnight in medium containing IL-2 (100 U/mL). The following morning TSA T were plated 5:1 and 10:1 with tumor cells that were stained with Cell Trace Dye. At the indicated time points, co-cultures were labeled with FcR block, dead cell exclusion dye and antibodies against CD3 and CD45. Tumor targets were identified as CellTraceViolet+CD3-/CD45- cells. Healthy cell targets were CellTrace+CD3+CD45+. Effector cells were identified as CellTrace-Violet-CD3+CD45+. Compensation was performed using single stained cells. Acquisition was performed on a Beckman Coulter CytoFlex S using CytExpert version 2.2.0.97 software. Analysis was performed using Flow Jo v 10.5. Representative dot plot from a 1:10 target:effector ratio co-culture showing flow stability gating, forward and side scatter cell identification, viable cells and targets+effectors.

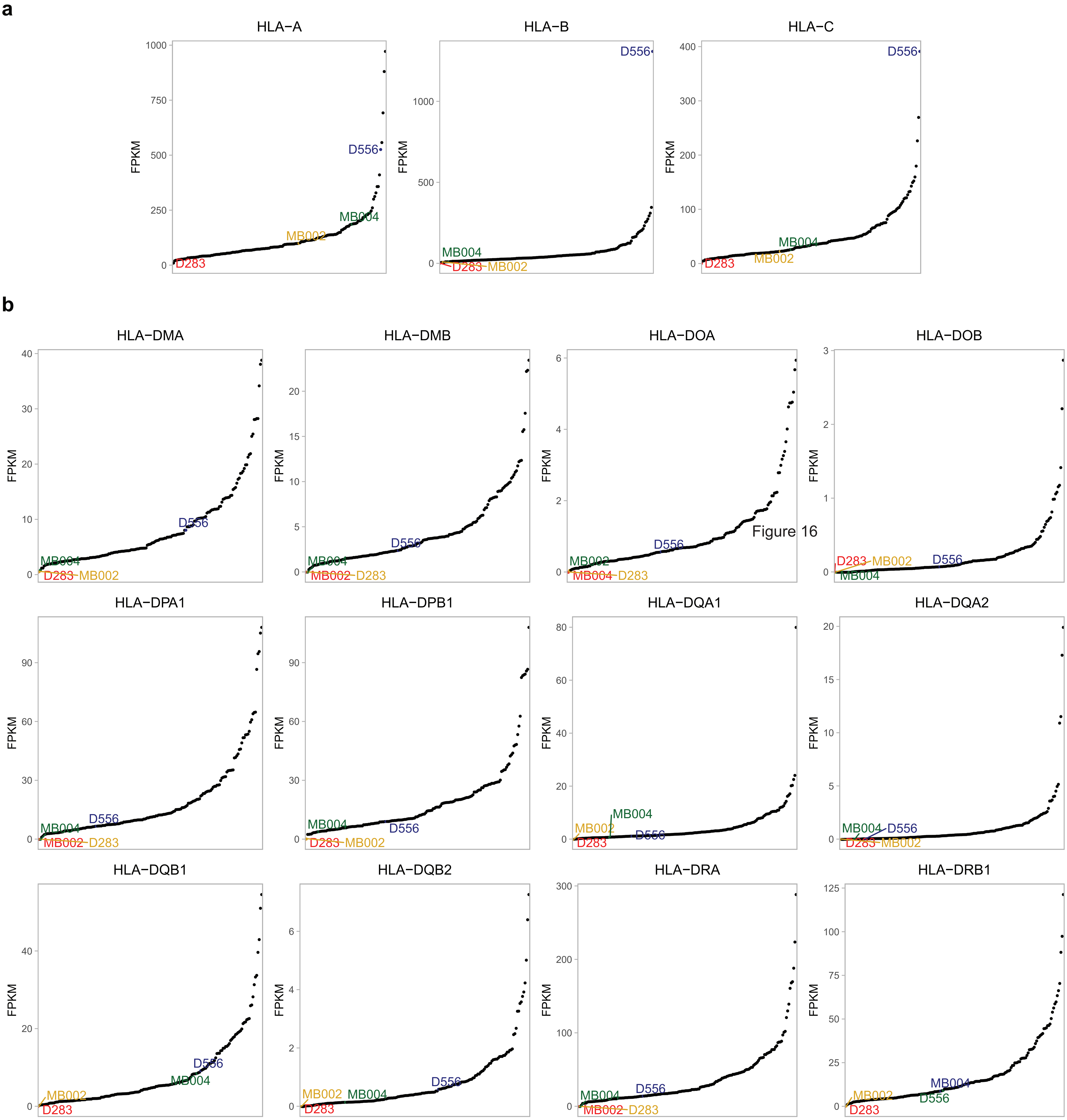


**Supplementary Fig. 19**

**Supplementary Fig. 19. HLA-I and HLA-II expression in medulloblastoma cell lines and primary tumors.** mRNA expression levels of HLA-I (**a**) and HLA-II (**b**) determined by RNA-seq. Fragment per kilobase and million reads (FPKM) are shown in the y-axis.

### SUPPLEMENTARY DATASET LEGENDS

Supplementary dataset 1. Clinical and tumor sample information.

Supplementary dataset 2. Summary of medulloblastoma tumor proteomic data.

Supplementary dataset 3. List of novel peptides detected in medulloblastoma tumors.

Supplementary dataset 4. Patient 7316-3778 peptides used for T cell stimulation

Supplementary dataset 5. Top 10 clonotypes within the 7316-3778 TSA T population

Supplementary dataset 6. Summary of medulloblastoma cell line proteomic data.

Supplementary dataset 7. List of novel peptides detected in medulloblastoma cell lines.

Supplementary dataset 8. HLA typing of MB002 and D556 cell lines and healthy donors used in this study.

Supplementary dataset 9. MB002 and D556 cell line peptides used for T cell stimulation.

Supplementary dataset 10. Antibodies used in this study.
