## Supplementary Data 5 for "Proteogenomic Discovery of Neoantigens Facilitates Personalized Multi-antigen Targeted T cell Immunotherapy for Brain Tumors"

**Supplementary dataset 5.** Top 10 clonotypes within the patient 7316-3778’s TSAT population

| **Clone** | **Productive**  **Frequency (%)** | **Amino Acid** | **V Resolved** | **D Resolved** | **J Resolved** |
| --- | --- | --- | --- | --- | --- |
| **1** | 18.03229239 | CSVHGGVSYEQYF | TCRBV29-01 | TCRBD02-01*01 | TCRBJ02-07*01 |
| **2** | 4.71198475 | CASTLGAGAYNEQFF | TCRBV27-01*01 | TCRBD02-01*02 | TCRBJ02-01*01 |
| **3** | 4.20283089 | CASSRVEPNEQFF | TCRBV12-03/12-04*01 | TCRBD02-01 | TCRBJ02-01*01 |
| **4** | 3.67230439 | CASSQDGGSYNEQFF | TCRBV03-01/03-02*01 | TCRBD02-01 | TCRBJ02-01*01 |
| **5** | 3.23060319 | CASRKRQDQPQHF | TCRBV12-03/12-04*01 | TCRBD01-01*01 | TCRBJ01-05*01 |
| **6** | 2.08739439 | CASRRVHSNQPQHF | TCRBV19-01*01 | TCRBD02-01*02 | TCRBJ01-05*01 |
| **7** | 2.08603018 | CASSLSRAGREQYF | TCRBV12 | TCRBD02-01*02 | TCRBJ02-07*01 |
| **8** | 2.04116279 | CASSSLGRTGVSEQYF | TCRBV07-08 | TCRBD01-01*01 | TCRBJ02-07*01 |
| **9** | 1.86063221 | CASRRRQDQPQHF | TCRBV12-03/12-04*01 | TCRBD01-01*01 | TCRBJ01-05*01 |
| **10** | 1.84183641 | CASSPRTSGRGETQYF | TCRBV05-01*01 | TCRBD02-01*02 | TCRBJ02-05*01 |
