## Supplementary Data 8 for "Proteogenomic Discovery of Neoantigens Facilitates Personalized Multi-antigen Targeted T cell Immunotherapy for Brain Tumors"

| **HLA** | **CELL LINE** | | **HEALTHY DONORS** | | | | |
| --- | --- | --- | --- | --- | --- | --- | --- |
|  | **D556** | **MB002** | **MB002**  **(1)** | **MB002**  **(2)** | **MB002**  **(3)** | **D556**  **(4)** | **D556**  **(5)** |
| A | 02:61/02:458; 23:01/23:54; | 30:11/30:02  01:01/01:215 | 03:01  23:01 | 01:01  68:02 | 02:01  23:01 | 02:01  23:01 | 01:01  68:02 |
| B | 44:257/44:185; 45:01 | 40:01  15:112/15:02 | 07:02  40:01 | 14:02  07:02 | 30:01  44:01 | 30:01  44:01 | 14:02  07:02 |
| C | 12:24/12:05; 06:87/06:27 | 07:02  08:72/08:01 | 03:04  07:02 | 07:02  08:02 | 07:02  04:01 | 07:02  04:01 | 07:02  08:02 |
| F | 01:01 | 01:01 |  |  |  |  |  |
| DPA1 | 01:03 |  |  |  |  |  |  |
| DPB1 | 04:02/16:01/  57:01 | 05:01/16:01/  38:01/22:01/63:01 | 04:01  04:01 | 02:01  04:01 | 02:01  04:01 | 02:01  04:01 | 02:01  04:01 |
| DQA1 | 01:01  02:01:01 |  |  |  |  |  |  |
| DQB1 | 05:01/05:03  03:03 | 06:01 | 03:01  06:02 | 03:01  06:02 | 02:02  05:01 | 02:02  05:01 | 03:01  06:02 |
| DRA | 01:01 |  |  |  |  |  |  |
| DRB1 | 07:01/07:01 | 15:02/15:01 | 11:03  15:01 | 13:01  15:01 | 01:01  07:01 | 01:01  07:01 | 13:01  15:01 |
| DRB3 | 02:02/02:01 |  | 02:02 |  |  |  |  |
| DRB4 | 01:01  01:03:01 |  |  |  | 01:01 | 01:01 |  |
| DRB5 |  | 01:01 | 01:01 | 01:01 |  |  | 01:01 |
| DRB9 | 01:01 |  |  |  |  |  |  |

**MB002 Healthy Donor 1**

Positive T cell response to pooled peptides 3-6 & 7-10 – abrogated in presence of anti-Class II;

donor DRB1*15:01 & DRB5*01:01 positive

**MB002 Healthy Donor 2**

Positive T cell response to peptides 6 and 7 – strong positive; donor C*07:02 DRB1*15:01 & DRB5*01:01 positive

**MB002 Healthy Donor 3**

Positive T cell response to peptides 6 and 7 – weak positive; donor C*07:02 positive, DRB1*15:01 & DRB5*01:01 negative

*Data support binding of peptides 6 and 7 through DRB1*15:01 &/or DRB5*01:01 alleles, consistent with CD4+ Flow data and anti-class II ELISPOT data.*

**D556 Healthy Donor 4**

Positive T cell response to peptides 6, 20

**D556 Healthy Donor 5**

Positive T cell response to peptide 14
